# Employing Machine Learning Techniques to Detect Protein-Protein Interaction: A Survey, Experimental, and Comparative Evaluations

**DOI:** 10.1101/2023.08.22.554321

**Authors:** Kamal Taha

## Abstract

This survey paper provides an in-depth analysis of various machine learning techniques and algorithms that are utilized in the detection of PPI (Protein-Protein Interactions). For every technique examined, the paper evaluates its efficiency, shortcomings, possibilities for enhancement, and outlook for the future. A major challenge in current survey papers focusing on machine learning algorithms for PPI identification is the successful categorization of these algorithms. To overcome this challenge, the paper introduces a novel hierarchical taxonomy that organizes algorithms into more intricate categories and distinct techniques. The proposed taxonomy is constructed on a four-tier structure, beginning with the broad methodology category, and ending with specific sub-techniques. This structure facilitates a more systematic and exhaustive categorization of algorithms, aiding researchers in grasping the connections between different algorithms and techniques. Included in the paper are both empirical and experimental assessments to classify the various techniques. The empirical assessment judges the techniques according to four standards. The experimental evaluations carry out the following rankings: (1) the algorithms that employ the same specific sub-technique, (2) the different sub-techniques that employ the same technique, (3) the different techniques that employ the same methodology sub-category, and (4) the different methodology sub-categories within the same methodology category. By merging the new methodological taxonomy, empirical analyses, and experimental evaluations, the paper provides a multifaceted and thorough comprehension of the machine learning methods and algorithms for PPI detection. This synthesis helps researchers make well-informed decisions. In its conclusion, the paper furnishes crucial insights into the future possibilities of machine learning techniques for PPI identification, underscoring potential advancements and areas ripe for continued exploration and development.

## 1. Introduction

Proteins are substantial molecular structures formed by extensive chains of amino acid residues, fulfilling various roles within living beings. These roles encompass activities like DNA replication, stimulus response, and molecule transportation. The well-orchestrated collaboration of protein molecules in Protein-Protein Interactions (PPIs) is essential in biological processes, and the overall health of an organism heavily depends on these interactions. Interruptions or irregularities in PPIs can lead to grave issues such as cervical leukemia, tuberculosis, and different neurological diseases. PPIs represent the physical contacts between two or more protein molecules, characterized by their precision. These contacts occur due to a biochemical incident, steered by forces like electrostatic power and hydrophobic effects, reflecting the molecular relationships within a living organism’s cellular or biological context.

Significantly contributing to the formation of macromolecular structures and enzyme complexes [1], PPIs are indeed central to nearly every biological process in living cells. Their influence stretches far beyond mere structural roles; they act as regulators and facilitators of life’s most complex tasks. Exploring these interactions is fundamental to understanding cell functions, providing a microscopic view into the inner workings of cells, tissues, and entire organisms. By uncovering disease mechanisms, PPIs shed light on the intricate pathways that lead to various health conditions, serving as both warning signals and potential targets for intervention. Identifying possible paths for novel treatments through PPIs is an ongoing pursuit that has the potential to revolutionize medical practice by offering therapies tailored to individual protein interactions. PPIs govern various biological functions, including cell interactions and the regulation of metabolism and growth [2], thereby controlling the vitality of life at the most fundamental level, and reinforcing the importance of their study in modern biology.

The accumulation of protein interaction data has led to the emergence of PPI Networks, displaying common traits with other networks, such as scale-free and small-world characteristics [3–7]. The study of PPI is of great importance and features in esteemed international journals and conferences like Nature [13], Science [8, 9], and Proceedings of the National Academy of Sciences [10, 11]. As the field advances, managing PPI data demands professional data support, with numerous comprehensive and regularly updated databases available to assist [12, 13]. As research advances, PPI data is beginning to display characteristics typical of big data. This necessitates expert management and support in utilizing the data for PPI studies. There is now a substantial number of open databases that are content-rich and frequently updated to assist in this area.

Protein interaction information is mainly gathered through diverse methods such as high-throughput screening technology [12, 13], predictive computation techniques for exploration [14–17], and literature searches [18, 19], among others. The amalgamation of these various sources helps to promote continuous research and innovation in the field. Several open databases, including the likes of BIND [20], DIP [21], IntAct [22], and HPRD [23], are compiled through the alignment of multiple processes. Also, open-source software applications have been developed around these databases, facilitating the extensive and unrestricted selection, storage, and querying of PPIs. As detection techniques and database technologies advance, so does the breadth and depth of protein interaction research.

Traditional methods of identifying PPI, such as Yeast Two-Hybrid screening or affinity purification, have their limitations. They are labor-intensive, costly, and restricted in scalability, often proving inadequate for the burgeoning complexity of modern biology. The surge in genomic and proteomic data demands innovative computational techniques to predict and interpret PPIs, thereby advancing avenues like drug discovery, disease modeling, and personalized medicine. Machine learning (ML), a branch of artificial intelligence, has emerged as a powerful tool in this context, showing remarkable proficiency in pattern recognition within complex data [7, 24, 25]. Using ML for PPI prediction is not just an exciting technological advance; it’s a hopeful and promising area that integrates various biological data types to produce insights previously unattainable through conventional methods. This application of ML in PPI prediction has the potential to radically transform our understanding of biological systems, reshaping our approach to biomedical science and paving the way for unprecedented advancements in healthcare and disease management.

Nevertheless, employing ML in PPI prediction is not without its hurdles, such as managing imbalanced datasets, selecting appropriate features, optimizing hyperparameters, and validating predictions. These challenges aren’t merely technical obstacles; they are central to the creation of models that are both robust and trustworthy. Addressing these issues requires a combination of scientific acumen, computational expertise, and a nuanced understanding of biological context. Meeting these challenges is not just a matter of refining the technology; it’s about crafting a toolset that’s suitable for the real-world complexity of biological research and practical implementation in medical and scientific applications.

The aim of this paper is to carry out an extensive analysis of contemporary and sophisticated ML algorithms for PPI detection through hands-on **empirical and experimental evaluations**. To fulfill this objective, we introduce a **classification system** rooted in methodology that organizes algorithms into nested hierarchical, specific, and meticulously detailed categories. This results in a more exact and finely-tuned categorization of various methods. In our analysis, we examined over 100 papers that encompass the finely-detailed and specific techniques related to ML PPI. For each method, we consulted reputable publishers and, to ensure that our paper selection is both current and represents the forefront of the field, we rated them based on their recency and innovation level, selecting the top ones that offered a broad spectrum of information about the technique. Our exhaustive evaluation is geared towards shedding light on the strong and weak points of the different ML algorithms applied to PPI, thus guiding subsequent research in this domain. This approach yields a thorough and intricate comprehension of today’s ML PPI algorithms and their practical applications.

### 1.1 Motivation and Key Contributions

Survey papers that focus on machine learning (ML) algorithms for predicting PPI currently grapple with the challenge of properly categorizing these algorithms. The present categorizations are too broad, lacking in particularity, and unable to differentiate between specific techniques. This vagueness can lead to misclassification of unrelated algorithms and inconsistent evaluations using the same criteria. This paper introduces a novel methodological taxonomy that organizes algorithms into finer categories and distinct techniques. With a four-tier structure, ranging from the broad methodology category to specific sub-techniques, this taxonomy enables a more organized and thorough classification of algorithms, helping researchers recognize connections between different algorithms and techniques.

The primary goal of this paper is to conduct an extensive survey of ML algorithms used in predicting PPI, employing a common sub-technique, technique, sub-category, and category. By leveraging this new taxonomy, researchers can more precisely compare and evaluate algorithms, deepening their understanding of each method’s pros and cons. Also, the taxonomy provides a structure for ongoing research, steering the creation and assessment of innovative algorithms. This paper significantly enriches the domain of ML-driven PPI detection by delivering a more all-encompassing and methodical way to categorize algorithms. It is anticipated that researchers will embrace this taxonomy, thus propelling the creation of more precise and efficient algorithms.

Not only does this survey introduce a thorough framework for classifying ML-based PPI prediction algorithms, but it also incorporates **empirical and experimental** assessments to gauge the efficacy of various methods. Our **empirical examination** scrutinizes techniques for ML-driven PPI detection using four criteria. Our **experimental evaluation** carry out the following rankings: (1) the algorithms that employ the same specific sub-technique, (2) the different sub-techniques that employ the same technique, (3) the different techniques that employ the same methodology sub-category, and (4) the different methodology sub-categories within the same methodology category.

The complete evaluation methodology equips researchers with the tools needed to discern fine distinctions between closely related algorithms and techniques. This allows them to select the approach best suited for their specific PPI prediction task using ML. The combination of a methodological taxonomy, empirical assessments, and experimental comparisons contributes to a more profound and nuanced understanding of the available algorithms for ML-based PPI identification. Researchers are thereby empowered to make educated choices in technique selection. This approach symbolizes a remarkable and meaningful step forward in PPI identification research, offering several essential advantages that streamline the process of algorithm selection.

### 1.2 Our Proposed Methodology-Based Taxonomy

We divide the algorithms utilizing ML to detect PPI into two primary categories: those based on Deep Learning and those based on traditional machine learning. Each of these categories is further broken down into four levels, with every subsequent level becoming more specific. Our hierarchy for methodology-based taxonomy is structured as follows: **methodology category → methodology sub-category → methodology techniques → methodology sub-techniques**. This hierarchical structuring allows us to pinpoint precise techniques or sub-techniques at the final stage. **Fig. 1** illustrates this methodology-based classification system. The benefits of our taxonomy are numerous, and they include:

1. *Improved Organization:* The taxonomy provides an orderly framework for displaying the survey’s findings. Through the classification of related methods, the hierarchical layout assists readers in understanding the paper’s logical progression.
2. *All-inclusive Examination:* The taxonomy guarantees a thorough examination by encompassing all pertinent methods. The hierarchical structure supports this coverage, assisting in identifying areas requiring further research and exploration.
3. *Facilitation of Technique Comparison:* Our classification system simplifies the comparison of different research techniques. By grouping like techniques and underlining their commonalities and differences, we can more easily discern the respective strengths and weaknesses of various approaches.
4. *Enhancement of Reproducibility:* By providing a clear techniques’ outline, our taxonomy enhances research reproducibility.

**Fig. 1:**
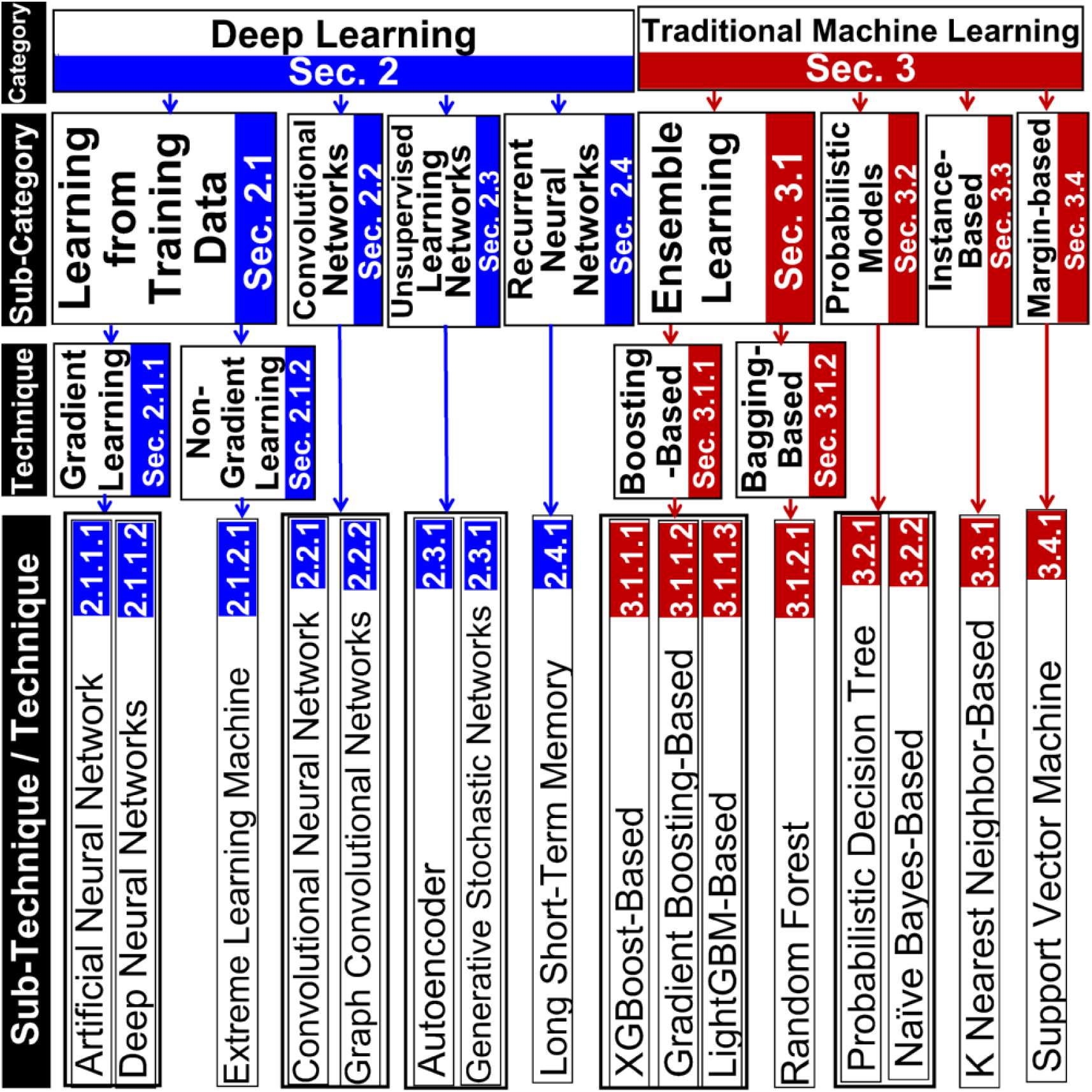
Our methodology-based taxonomy that categorizes the machine learning algorithms for the detection of PPI into fine-grained classes in a hierarchical manner, as follows: methodology category → methodology sub-category → methodology technique → methodology sub-technique. For each category, sub-category, technique, and sub-technique, the figure also shows the section number in the manuscript that discusses it.

### 1.3 Methodology for Selecting Papers

The process we followed for selecting and analyzing the papers to be incorporated into this survey paper is detailed as follows:

*Keywords and Search Criteria*: The specific keywords and phrases that we used to guide the literature search include the following: “deep learning in PPI detection,” “traditional machine learning in PPI detection,” “protein-protein interaction,” “machine learning,” etc. We utilized Boolean operators to refine the search.

1. *Databases and Sources:* We identified the databases and sources that will search, which included the following: PubMed, IEEE Xplore, ACM, Elsevier, and Springer.
2. The criteria that papers must meet to be included in the survey includes the following:

- Published in a peer-reviewed journal or conference.
- Specific to machine learning techniques applied to PPI detection.
- Meeting a certain threshold of impact.
3. *Screening Titles and Abstracts:* We carried out an exhaustive examination of the titles and abstracts of the papers we initially gathered, considering our predefined inclusion and exclusion criteria. Papers that were clearly misaligned with our research goals were specifically omitted from consideration.
4. *Full Paper Evaluation:* We examined the entire collection of selected papers, meticulously assessing the content, methodology, research contributions, and relevance of each paper to ensure alignment with our study’s focus.

## 2. Deep Learning-Based Category

> The use of deep learning methods to predict protein-protein interactions (PPIs) is a vital field of study that has considerable consequences in both biology and medicine. The procedure begins with gathering information about recognized PPIs and extracting characteristics from amino acid sequences during the data collection and preprocessing stages. Various deep learning models, including Convolutional Neural Networks (CNNs), Recurrent Neural Networks (RNNs), and graph-based structures, are utilized in this process. Each of these models can understand different elements of protein structures and sequences. For instance, CNNs analyze spatial characteristics, RNNs handle extensive sequences, and graph-based models depict the topological attributes of proteins. By adding extra features through feature engineering, the models’ ability to predict can be strengthened, as it includes specialized knowledge in the field. The models are then trained and validated on a subset of known PPIs, and hyperparameters are fine-tuned to prevent overfitting. The procedure of deep learning is illustrated in Fig. 2.

**Fig. 2:**
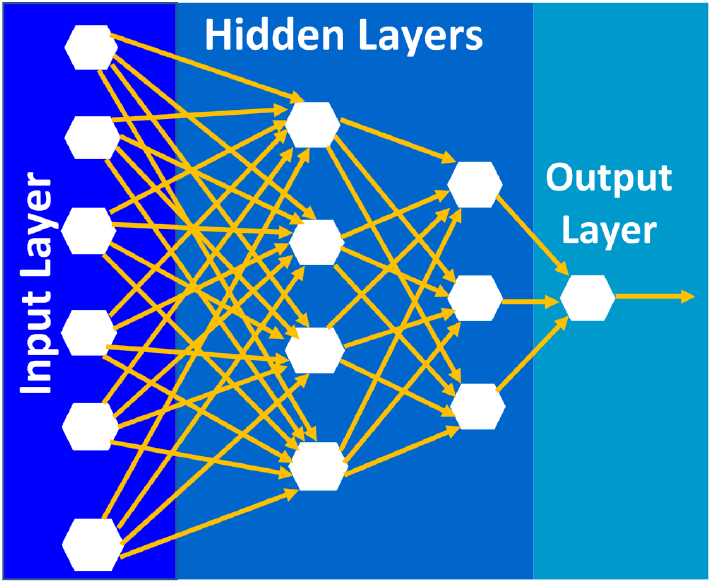
The procedure of deep learning is illustrated in the figure.

### 2.1 Learning from Training Sub-Category

#### 2.1.1 Gradient Learning Technique

##### 2.1.1.1 Artificial Neural Network (ANN) Sub-Technique

> Utilizing Artificial Neural Networks (ANNs) to predict PPI are computing systems inspired by the biological neural networks of animal brains. They are composed of interconnected artificial neurons or nodes, where information is processed as it passes through the network. ANNs can be shallow, with only one or two hidden layers, or deep, with many hidden layers. By analyzing properties such as amino acid sequences, ANNs can build predictive models of these interactions. This technique provides a faster and more cost-effective alternative to traditional methods of studying PPI, having applications in areas like drug development and disease diagnosis. It represents a significant advancement in the field, although its accuracy relies on the quality of training data and the chosen algorithms, necessitating continuous research and refinement.

Vyas et al. [26], showcased a novel method for predicting disease-related protein-protein interactions (PPIs) by employing a Genetic Programming (GP) based Symbolic Regression (SR) technique integrated with an Artificial Neural Network (ANN). In their case study, they used a dataset comprising 135 PPI complexes associated with cancer. This information allowed them to create a generic model for predicting PPIs, exhibiting both high accuracy in predictions and a robust ability to generalize. Bakar et al. [27], crafted a method using ANN to PPIs by analyzing secondary structures, co-localization, and function annotation. This approach utilizes hybrid machine learning algorithms to foresee the interactions of proteins specifically within yeast. Goodacre et al. [28] introduced an ANN-based approach to identify new non-synonymous Single Nucleotide Polymorphisms (nsSNPs) that could potentially disrupt the interaction between HIV-1 and human proteins. The model was trained using machine learning, employing physical and energy scores derived from HADDOCK docking. To differentiate between variants that have either completely or partially lost binding ability, a random forest classifier was utilized. This classifier was specifically designed to analyze the differences in docking scores between 166 mutant pairs and their corresponding wild-type forms.

##### 2.1.1.2 Deep Neural Networks (DNN) Sub-Technique

> Utilizing Deep Neural Networks (DNNs) to predict PPI involves the application of advanced deep learning algorithms to model and predict how proteins interact with each other. DNNs are a specific type of ANN that contains multiple hidden layers of nodes between the input and output layers. These additional hidden layers enable the network to model more complex and abstract features. DNNs are often employed in deep learning, where they can be trained to recognize patterns and make predictions based on large datasets. The technique can also integrate different biological attributes such as sequence and structural information, making it highly versatile.

Li and Yu [29] introduced a deep neural network that combines convolutional and recurrent layers. They implemented a feature embedding layer to convert a sequence into a more compact matrix form. The model includes several convolutional neural network layers, as well as stacked bidirectional relational neural network layers, designed to understand both local and global context information from the compressed matrix. To construct the classifier for the prediction task, fully connected layers and softmax layers were added to the top of the model. Li et al. [30] introduced a self-attention-based deep learning neural network approach for predicting PPIs. Initially, the protein sequence information is encoded using AAC, CT, and AC. Then, the deep neural network is merged with a self-attention method to efficiently extract features. This process adjusts the weight of the sequence to highlight essential characteristics, thereby creating a network model that can comprehensively extract information from the protein sequence. Wang et al. [31] presented a DNN method for improving the prediction performance of protein interaction relationship pairs.

By adjusting factors like the learning rate and using different parameters, including the activation function, they were able to enhance the model’s ability to predict these interactions. Tran et al. [32] suggested a method that integrates deep learning with feature fusion for PPI prediction, merging handcrafted features with protein sequence embedding to represent protein sequences. In this approach, the authors employ a fully connected layer to understand the nonlinear relationships of the inputs and to reduce the data from a high-dimensional space to a lower-dimensional one. The feature combination module is composed of a merge layer, a fully connected layer, a batch normalization layer, and a dropout layer. Ultimately, a feature vector representing the input protein pairs is created and utilized by the classification module to predict interactions.

#### 2.1.2 Non-Gradient Learning Technique

##### 2.1.2.1 Extreme Learning Machine (ELM) Sub-Technique

> Utilizing Extreme Learning Machine (ELM) to predict PPI is an approach that combines computational power with biological research. ELM is a type of feedforward neural network with a single hidden layer, and it’s known for its rapid training process. Unlike traditional neural networks, the weights connecting the input layer to the hidden layer are randomly assigned and do not change during training. Only the weights connecting the hidden layer to the output layer are tuned, allowing for faster learning. This is especially beneficial in applications that require quick and efficient modeling of complex relationships.

Deddy et al. [33] introduced a quick prediction algorithm for PPI that utilizes the ELM algorithm. By fully leveraging the ELM algorithm, they were able to achieve favorable results in protein prediction. You et al. [34] introduced a technique for PPI prediction that relies solely on the information from protein sequences. This method was fashioned using the ELM learning algorithm, coupled with a depiction of local protein sequence descriptors. These local descriptors consider the interactions between residues in both continuous and discontinuous areas of a protein sequence. Consequently, this approach allows for the extraction of additional PPI information from the protein sequences. Sikandar et al. [35] have examined new computational techniques for disease-gene association, utilizing advanced biological and topological characteristics. This method achieves an understanding of protein-protein interaction (PPI) within the human genome and facilitates the discovery of hereditary gene-disease relationships through topological features drawn from the PPI network. During the training process, the back-propagation approach is employed in the ELM model, allowing data to flow back through the network. The network’s weights remain fixed during the validation phase, wherein the trained model is imported to estimate the real data. The ELM model comprises an input layer, several hidden layers, and a single output layer.

You et al. [36] introduced a framework known as Low-rank Approximation Kernel LELM for the automatic detection of human PPI from primary protein sequences. This framework consists of three main phases: 1) converting each protein sequence into a matrix constructed from various adjacent amino acids; 2) using the low-rank approximation model to address the matrix, finding its most minimal rank representation; and 3) employing a robust kernel extreme learning machine to estimate the likelihood of PPI. Qiu et al. [37] introduced a method to predict hot spots at the interfaces of PPIs, utilizing the ELM. The selection classifier in this method consists of two parts: the first part targets the removal of unnecessary features from the original ones without affecting the prediction, while the second part focuses on building the final prediction model based on that prediction. You et al. [38] introduced a sequence-based approach to identify PPIs by utilizing the ELM in conjunction with a unique representation employing auto covariance (AC). This method properly considers the neighboring effect, allowing for the extraction of more PPI information from the protein sequences.

Huang [39] introduced a method for predicting PPIs that relies solely on the information derived from protein sequences. This approach is formulated using the ELM in conjunction with Chou’s Pseudo-Amino Acid Composition. The ELM classifier has been chosen as the prediction engine for this methodology. Li et al. [40] introduced a method for predicting PPIs using protein sequences. Initially, the protein sequences are converted into a Position Weight Matrix (PWM), and the Scale-Invariant Feature Transform algorithm is utilized to extract features. Subsequently, Principal Component Analysis (PCA) is applied to reduce the dimensionality of the features. Finally, a Weighted ELM classifier is used to make predictions about PPIs.

### 2.2 Convolutional Networks Sub-Category

#### 2.2.1 Convolutional Neural Network (CNN) Technique

> Utilizing Convolutional Neural Networks (CNNs) to PPIs is a sophisticated technique that combines elements of machine learning and bioinformatics. This approach focuses on applying the unique ability of CNNs to analyze and recognize spatial patterns in data to the complex structures and sequences that define proteins. By representing proteins as sequences or structural data, CNNs can extract essential features and relationships through the application of convolutional and pooling layers. Subsequent fully connected layers interpret these patterns to predict how pairs of proteins might interact, a key insight for understanding biological functions, disease mechanisms, and drug discovery. Despite its promising contributions, this method faces challenges such as the complexity of protein structures, the immense number of possible interactions, and the need for significant computing resources.

Cai and Zhu [41] introduced a technique founded on particle swarm optimization (PSO), enabling the automatic construction of a deep convolutional neural network (CNN) for the identification of essential proteins. Utilizing a NAS algorithm that relies on particle swarm optimization, the authors crafted the CNN. This approach results in the acquisition of low-dimensional dense vectors, which encompass intricate topological features from PPI networks. Zhang et al. [42] introduced a method for PPI extraction that utilizes the residual CNN. By stacking additional convolutional modules through residual connections, this model is designed.

It diminishes the need for conventional biomedical natural language processing tools like the dependency parser. Yuan et al. [43] introduced a predictor for PPI prediction, using deep transfer learning. They developed a deep PPI detector to forecast unknown PPIs, thereby completing the known PPI network. The system is composed of a transfer CNN feature extractor, which is pre-trained by GO (Gene Ontology) annotation, and a semi-supervised SVM classifier. Dutta et al. [44] introduced a four-stage ensemble clustering method for detecting PPI and gene clustering. This approach utilizes a multi-layer perceptron and a deep CNN to generate a consensus partitioning.

#### 2.2.2 Graph Convolutional Networks Technique

> Utilizing Graph Convolutional Networks (GCNs) to predict PPI is an advanced approach in computational biology that employs GCNs to model and understand the complex interactions between proteins. GCNs can capture the intricate spatial relationships found in the three-dimensional structures of proteins, considering both their local and global characteristics. By considering the proteins’ three-dimensional shapes, amino acid sequences, and other biological attributes, GCNs can learn to identify patterns that enable the prediction of unknown interactions.

Li et al. [45] suggested an approach to identify PPI by incorporating residual connections, dense connections, and dilated convolutions into deep Graph Convolutional Networks (GCNs) architectures. Specifically, they employed these concepts, which had previously proved effective in the training of deep Convolutional Neural Networks (CNNs), including the use of residual connections, dense connections, and dilated convolutions. Voytetskiy et al. [46] carried out an implementation of a GCN approach for PPI. They experimented with three distinct GNN architectures: GCN, Graph Attention Network (GAT), and an inductive representation learning network. In their testing, the GNN models demonstrated superior performance compared to the state-of-the-art Recurrent Neural Network (RNN) model.

Zhu et al. [47] introduced a semi-supervised network embedding model aimed at enhancing PPI and protein complex detection. They utilized graph convolutional networks to efficiently identify densely connected subgraphs. Specifically, the authors crafted a three-layer GCN to deeply analyze the structure of PPI networks, ensuring the preservation of the second-order proximity. Jha et al. [48] employed Graph Convolutional Network (GCN) and Graph Attention Network (GAT) to predict PPI, leveraging the structural information and sequence features of proteins. They constructed protein graphs using their PDB files, containing the 3D coordinates of atoms. These protein graphs symbolize the amino acid network, or the residue contact network, where each node corresponds to a residue. A connection between two nodes is established if a pair of atoms (one from each node) is found within a specified threshold distance.

### 2.3 Unsupervised Learning Networks Sub-Category

#### 2.3.1 Autoencoder Technique

> Utilizing an autoencoder to predict PPI is a method that involves compressing data into a lower-dimensional form and then reconstructing it, learning the essential features that capture the underlying structure of the data. Autoencoders can be trained on known interactions to identify the key features that define PPI relationships. This learning process often involves encoding the amino acid sequences or the three-dimensional structure of the proteins into a continuous latent space, where patterns of interaction can be detected. The compressed representations can capture subtle correlations between proteins, which may be invisible through other methods. After training, the autoencoder can be used to predict interactions between unknown proteins, or to predict new interactions between known proteins.

Atashin et al. [49] introduced a technique to address the PPI prediction issue by utilizing a denoising autoencoder. This method employs a Denoising Auto-Encoder to learn robust characteristics. These learned features are then applied to train a multilayer feedforward neural network, enhancing the system’s overall performance. Sharma and Singh [50] introduced a method for accurate PPI prediction that makes use of an Autoencoder. This technique efficiently creates lower-dimensional, discriminative, and noise-free features. By incorporating both conjoint triad (CT) features and Composition-Transition-Distribution (CTD) features into the model, the authors were able to achieve promising results. Albu [51] unveiled a two-stage, sequence-oriented method for PPI prediction, which relies on supervised autoencoders. This method begins with the training of a denoising autoencoder that focuses on protein sequences. Subsequently, there is a supervised training phase where the model is taught to simultaneously predict if two proteins will interact and to rebuild the paired proteins involved.

Cao et al. [52] introduced a method for multi-network embedding that utilizes a deep variational autoencoder (VAE). This method employs the variational autoencoder to distill low-dimensional characteristics of proteins from several different interactive network datasets. Following this extraction, an SVM classifier is trained to predict PPI. Xiao et al. [53] introduced a PPI model founded on two core concepts: a) using a variational autoencoder model predicated on a Gaussian distribution as a fundamental PPI predictor to handle the sparse input PPI Network; b) integrating this basic PPI predictor into a flexible architecture, utilizing a strategy that relies on both block structures and probability summation. Luo et al. [54] introduced a PPI prediction algorithm that merges the sequence and network data of proteins using a variational graph autoencoder. Initially, they implemented various techniques to distill the characteristics of proteins from both their sequence and network details, subsequently compacting these features through principal component analysis. The authors also created a scoring function to evaluate the higher-order connectivity between proteins, resulting in a higher-order adjacency matrix.

Jha et al. [55] applied a stacked auto-encoder to address the problem of PPIs. Their model operates on a 92-length feature vector that represents an amalgamation of features drawn from the protein sequence through various methods. This vector includes evolutionary features derived from the PSI-BLAST algorithm, forecasted structural attributes, and seven specific physicochemical properties of amino acids. Sun et al. [56] utilized a Stacked Autoencoder (SAE) to examine sequence-based predictions of human PPIs. They found that models employing protein sequence autocovariance coding yielded the most promising results in both 10-fold cross-validation and when forecasting hold-out test sets.

#### 2.3.2 Generative Stochastic Networks Technique

> Generative Stochastic Networks (GSNs) are a class of generative models that aim to learn the underlying probability distribution of a dataset. GSNs typically consist of a series of hidden layers that transform random noise into data that resembles the distribution of the training set. They work by iteratively refining this noise through the layers, improving the resemblance at each step.

Zhang et al. [57] created a Generative Stochastic Model that incorporates both Functional and Topological Properties to delineate the formation processes of the PPI network and the associated functional profile. This model serves as a mechanism to grasp the interaction structures and the functional behaviors of proteins. By fusing functional and topological characteristics, they shaped the problem of identifying protein complexes in such a way that it focuses on detecting groups of proteins that often interact within the PPI network and exhibit similar annotation patterns in the functional profile. Through the concept of link communities, their method naturally accommodates overlaps among these complexes. Linder et al. [58] proposed an approach utilizing Generative Stochastic Networks, where the most prominent sequence positions are detected using learned input masks. Scramblers are trained to create Position-Specific Scoring Matrices (PSSMs), and they are used to interpret the consequences of genetic variants. Additionally, they help reveal non-linear relationships between cis-regulatory elements, explain binding septicity in PPI, and find the structural factors that determine the properties of de novo designed proteins.

Schweiger et al. [59] introduced a fresh view on PPI network modeling. They differentiate between two types of models: one that follows the duplication–divergence (DD) process, characterized by copying neighbors, and the other that doesn’t copy neighbors, with the Barabási–Albert (BA) preferential attachment model standing as a prime illustration of the latter. Wang et al. [60] built a dataset focusing on drug-likeness in PPI and introduced a deep molecular generative framework to create new molecules resembling drugs from the characteristics of the initial seed compounds. Drawing inspiration from previously published molecular generative models, this framework employs the essential features related to PPI inhibitors as input. It then advances deep molecular generative models for the original design of PPI inhibitors.

Wang et al. [61] introduced a prediction algorithm for PPI networks, utilizing Generative Stochastic Networks. This algorithm simulates the creation of a PPI network, effectively capturing underlying community structures. It then adopts an inference process that further refines the distribution of proteins’ memberships across different complexes. Subsequently, a specially designed distance measure calculates the similarity between two proteins based on the probabilities of them being in the same complex, thereby determining whether or not they interact with each other. Zhou and Troyanskaya [62] introduced a generative stochastic networks (GSN) model for PPI to learn the generative model of data distribution without the need to clearly define a probabilistic graphical model. Specifically, they applied this supervised extension of GSN by training a Markov chain to sample from a conditional distribution. This approach was used for the task of predicting protein structure.

### 2.4 Recurrent Neural Networks Sub-Category

#### 2.4.1 Long Short-Term Memory (LSTM) Technique

> Long Short-Term Memory (LSTM) is a specialized form of recurrent neural network that excels at recognizing long-range dependencies in sequential information. When applied to predicting PPI, proteins are characterized as sequences of amino acids and then transformed into a numerical format. This numerical encoding is used as input for the LSTM network, leveraging its ability to remember long-range dependencies. By training on data from known interactions, the LSTM network can discern the intricate patterns and relationships that dictate how proteins interact with each other.

Lu et al. [63] introduced a deep learning approach that utilizes a Bi-LSTM model to forecast PPI. This method automatically translates amino acids, representing the protein sequences, and then proceeds to extract the sequence characteristics for classification purposes. Yadav et al. [64] introduced a technique using deep B-LSTM that harnesses word sequences and dependency path-related information to detect PPI information from textual data. This model unifies the joint modeling of proteins and relations into one cohesive framework. Zeng et al. [65] introduced a deep learning framework designed to autonomously learn biological features without the need for pre-existing knowledge. By employing the node2vec technique, it can learn a more comprehensive representation of PPI network topologies than what a score function might provide. Bi-LSTM cells are used within the framework to identify non-local relationships in gene expression data.

Tsukiyama et al. [66] created an LSTM model in conjunction with word2vec to forecast PPIs between humans and viruses solely through the utilization of amino acid sequences. The model is trained on data with a significantly imbalanced ratio of positive to negative samples. Within this framework, word2vec retains the information regarding the patterns of local amino acid residues, while the LSTM goes further to learn the patterns of amino acids across the entire sequence context. Alakus and Turkoglu [67] suggested utilizing a deep learning method to forecast PPI. This approach initially identifies the groups of proteins. Next, it translates the protein sequences into numerical values, applying both protein signature techniques and ProtVec methods. In the final step, an LSTM model is employed to categorize and predict the interactions between the proteins.

## 3. Traditional Machine Learning Category

### 3.1 Ensemble Learning Sub-Category

#### 3.1.1 Boosting-Based Technique

##### 3.1.1.1 eXtreme Gradient Boosting (XGBoost)-Based Sub-Technique

> Utilizing XGBoost (Extreme Gradient Boosting), a renowned decision tree-based ensemble learning algorithm, enhances both accuracy and efficiency in predicting PPI. This approach entails the extraction of pertinent features from protein sequences, such as structural and physical attributes. The data is organized by partitioning it into training and validation sets, and the XGBoost algorithm is subsequently trained using this labeled interaction information. To further refine the model’s efficacy, fine-tuning of hyperparameters is often carried out, aiming to optimize predictive performance.

Chen et al. [68] introduced a two-layer model to predict host-pathogen PPI, addressing the difficulties related to the algorithms for feature representation and the imbalance in the data. This two-layer model is made up of two vital modules: XGBoost, utilized to diminish the data’s imbalanced ratio, and SVM, implemented to enhance performance. Additionally, SMOTE technology is integrated within the model as a critical element to mitigate the skewed ratio caused by the imbalanced data. Wang et al. [69] introduced a method for predicting PPI sites based on XGBoost. Initially, characteristic information about the protein is extracted using the pseudo-position specific scoring matrix, pseudo-amino acid composition, and hydropathy index. To address class imbalance, the synthetic minority oversampling technique is employed, and the kernel principal component analysis is applied to eliminate redundant characteristics. These optimized features are input into the XGBoost classifier to accurately identify PPI sites.

Beltran et al. [70] demonstrated that XGBoost surpasses SVM and Random Forest in predicting PPI, achieving better results in aspects such as accuracy, specificity, and the Matthews correlation coefficient. Mahapatra et al. [71] introduced a hybrid method that integrates a DNN with XGBoost for predicting PPI. This combined classifier utilizes a blend of three sequence-based features as inputs: amino acid composition, conjoint triad composition, and local descriptor. Within this system, the DNN uncovers concealed information by abstracting it layer-by-layer from the raw features, which are then processed by the XGBoost classifier. Zhong et al. [72] introduced a framework for predicting essential proteins that utilizes XGBoost models and includes multivariate features. Within this framework, the SUB-EXPAND-SHRINK method is applied for feature engineering. This technique helps in crafting composite features from the original ones and allows for the selection of a more optimal subset of features, enhancing the prediction of essential proteins. Wang et al. [73] introduced a method for predicting PPI sites based on XGBoost. Initially, characteristic information about the protein is extracted using the pseudo-position specific scoring matrix (PsePSSM), pseudo-amino acid composition (PseAAC), hydropathy index, and solvent accessible surface area (ASA) within a sliding window framework. To address class imbalance, the synthetic minority oversampling technique (SMOTE) is employed. Finally, these optimized features are input into the XGBoost classifier to accurately identify PPI sites.

##### 3.1.1.2 Gradient Boosting (G-Boosting) Sub-Technique

> Gradient Boosting (G-Boosting) builds a predictive model in a stage-wise fashion from a set of weak learners, typically decision trees. G-Boosting iteratively adds trees, usually shallow ones, to the model. In each iteration, a new tree is fit to the residual errors (the difference between the predicted and actual values) of the existing ensemble of trees. When applied to PPI, relevant features are extracted from protein sequences, such as their structural and physical characteristics. The data is then divided into training and validation sets, and the G-Boosting algorithm is applied using this labeled interaction information. The model may be further refined through the fine-tuning of hyperparameters to enhance performance. Fig. 3 shows the procedure of G-Boosting.

**Fig. 3:**
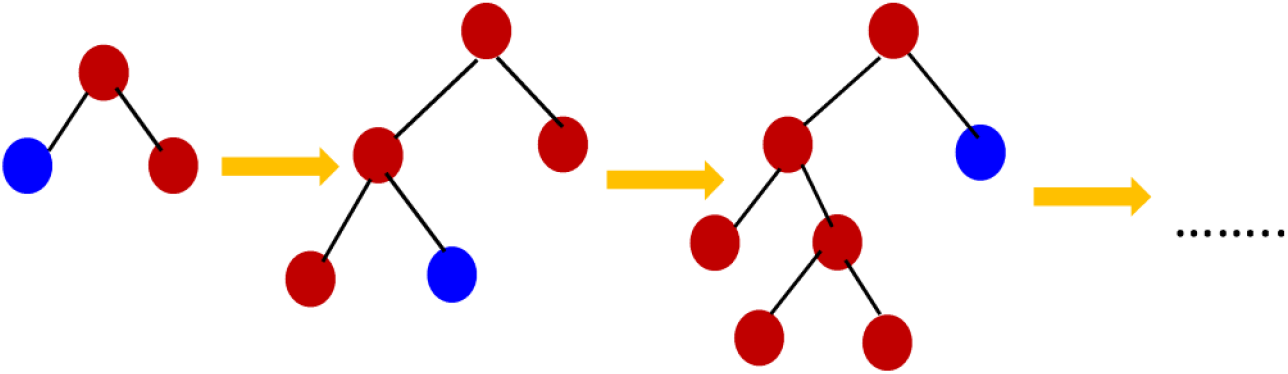
The procedure of Gradient Boosting is illustrated in the figure.

Lin et al. [74] presented the ensemble learning algorithm: boosting, as well as gradient boosting (G-boosting), paired with a couple of feature selection approaches. They designed boosting and G-boosting models utilizing structure features with the purpose of classifying hot spots and non-hot spots, along with Date-hub protein interfaces and Party-hub protein interfaces. Zeng et al. [75] introduced an ensemble machine learning model designed specifically to enhance the accuracy of essential protein prediction using just protein sequences. The model employs a composite structure, integrating multiple Gradient Boosting base classifiers. Furthermore, to mitigate the impact of imbalanced datasets, it incorporates a sampling technique. Mahapatra et al. [76] suggested a hybrid methodology that merges DNN and Extreme Gradient Boosting classifier (XGB) to predict PPI. This hybrid classifier utilizes a combination of three sequence-based characteristics, namely, amino acid composition, conjoint triad composition (CT), and local descriptor (LD) as input. The DNN operates to extract hidden information via a layer-by-layer abstraction from the raw features, which are subsequently passed through the XGB classifier.

##### 3.1.1.3 Light Gradient Boosting Machine (LightGBM) Sub-Technique

> LightGBM (Light Gradient Boosting Machine) works by constructing an ensemble of decision trees in an iterative manner, where each tree is built to correct the errors of its predecessors. It combines several unique techniques and optimizations to achieve efficiency, speed, and accuracy. LightGBM employs a histogram-based technique. This means that continuous values of features are divided into discrete bins, creating a histogram. This significantly speeds up the training process and reduces memory consumption. LightGBM adopts a leaf-wise growth strategy. It selects the leaf with the maximum delta loss to grow, allowing for more asymmetrical trees that can reduce more loss. While this may cause over-fitting on small datasets, it’s often controlled by other parameters such as “max_depth.”

Chen et al. [77] have put forward a method for the prediction of PPIs. In the first stage, they use pseudo amino acid composition, autocorrelation descriptor, local descriptor, and conjoint triad to pull out essential feature data. In the following step, an elastic net technique is employed to pinpoint the best feature subset and remove any superfluous features. In the final stage, LightGBM is utilized as the predictive model for determining PPIs. Zhan et al. [78] introduced a predictive model that utilizes both sequential and evolutionary data to forecast PPI interactions, as well as ncRNA and protein interactions. LightGBM is utilized as the consolidated classifier. For the prediction of ncRNA and protein interactions, a computational model is put to use. To derive the distinguishing characteristics from protein and ncRNA sequences, the pseudo-Zernike Moments and singular value decomposition algorithm are employed.

Hossain and Islam [24] introduced a method for identifying crucial proteins by employing a combination of a metaheuristic algorithm and a machine learning technique. They utilized three distinct classifiers within the objective function of their proposed strategy: LightGBM, Random Forest, and XGBoost. These classifiers were applied independently within the objective function to accomplish precision based on the molecular feature subset from the yeast dataset. The researchers further evaluated the performance by integrating an ensemble method with these individual classifiers. They utilized the LightGBM classifier with its standard parameter values.

#### 3.1.2 Bagging-Based Technique

##### 3.1.2.1 Random Forest Sub-Technique

> Random Forest to predict Protein-Protein Interaction (PPI) consists of multiple decision trees that work collectively to produce more accurate and robust predictions than a single decision tree. The Random Forest algorithm trains multiple decision trees on different subsets of the training data, each tree being built on a random sample of features. By doing so, Random Forest minimizes overfitting, which is a common challenge in machine learning. The predictions from individual trees are then aggregated, typically through a majority vote, to produce the final prediction. Hyperparameter tuning, such as adjusting the number of trees in the forest or the maximum depth of each tree, may be performed to enhance the model’s performance.

Wang et al. [79] introduced a framework to predict disease genes for hepatoma. This framework begins by extracting gene features from both the PPI network and gene ontology annotations using graph embedding techniques. Once these low-dimensional gene feature vectors are obtained, three traditional classifiers are applied to forecast the hepatoma-related genes: Random Forest (RF), SVM, and Logistic Regression (LR). Le and Kha [80] introduced a framework that employs a random forest algorithm and the FASTA sequence of amino acids for predicting PPI (Protein-Protein Interaction) networks. This method extracts the features of individual proteins and uses a neural network model to predict PPI directly, based on the attributes of the proteins.

Zhao et al. [81] introduced a new computational approach that includes a learning process to discover previously unidentified TSGs (Tumor Suppressor Genes). This method draws upon features extracted from PPI networks through an advanced network embedding algorithm. Multiple RF (Random Forest) models were developed, capable of discerning crucial differences between TSGs and other genes. Specifically, thirty random forest models were assembled and employed to predict hidden or latent TSGs. Khatun et al. [82] introduced a method to identify PPI using a machine learning approach. They trained a random forest classifier with autocorrelation features to construct the prediction model. The method incorporates autocorrelation (AtC) along with a supervised statistical learning framework, specifically utilizing random forest.

### 3.2 Probabilistic Model Sub-Category

#### 3.2.1 Probabilistic Decision Tree Technique

> A Probabilistic Decision Tree (PDT) is a statistical model that combines the principles of decision trees with probability theory. A PDT allows for uncertainty and probabilistic outcomes at each decision node. The decision at each node is based on probabilities. Training involves learning the probabilities at each decision node based on the observed frequencies in the training data. It might also involve learning the structure of the tree itself. Fig. 4 shows the procedure of decision tree-based classifier.

**Fig. 4:**
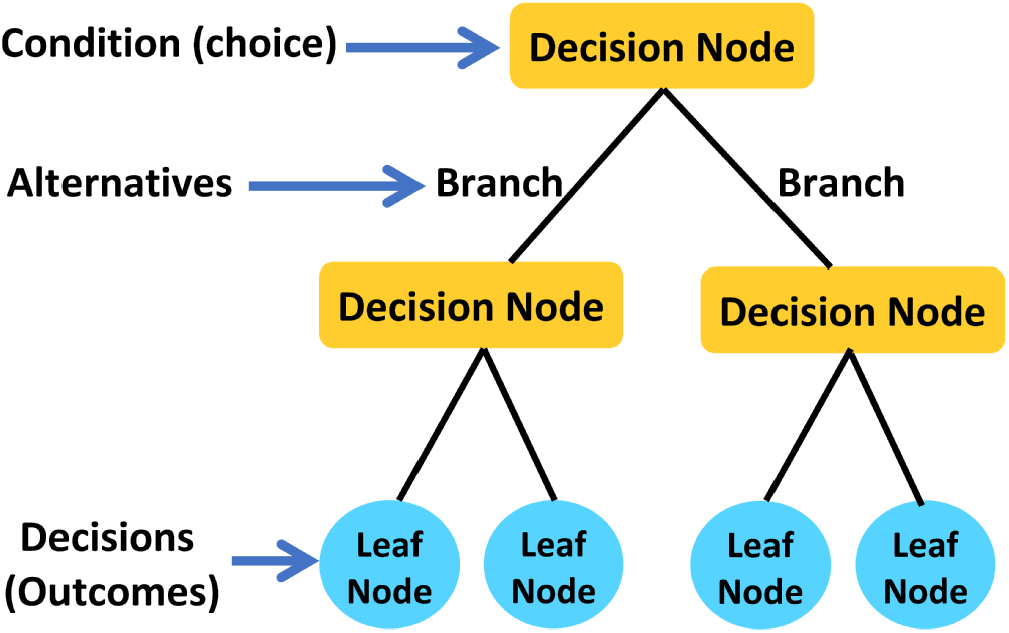
The procedure of Decision Tree-based classifier is illustrated in the figure.

Cecchini et al. [25] introduced a proficient method for predicting metabolic disease genes, addressing the challenge of imbalanced data. This method fuses PPI and miRNA-target interactions to build integrated networks. The topological properties of these networks are utilized as features for training machine learning classifiers. For the specific task of binary classification, a Probabilistic Decision Tree was employed. The concept of boosting was applied, which involves training an ensemble of weak learners using decision trees. Zaki and Alashwal [83] outlined a method to identify missing interactome links in the PPI network, thereby enhancing the detection of protein complexes. They located the missing links by extracting various topological features, which were then used alongside a two-class boosted decision-tree classifier to build a machine-learning model. This model is adept at differentiating between existing and non-existing interactome links. It was trained on a PPI network composed of 1,622 proteins and 9,074 interactions.

Yang et al. [84] applied a Probabilistic Decision Tree to forecast PPI scores based on proteins’ CETSA (Cellular Thermal Shift Assay) features. For a limited set of protein pairs where the predicted PPI scores did not align with the actual data, the authors employed an iterative clustering strategy to progressively decrease the count of these mismatched pairs. After completing the iterative clustering, the remaining protein pairs, which may possess atypical properties, were deemed scientifically valuable for additional biological study.

#### 3.2.2 Naïve Bayes Technique

> Applying a Naïve Bayes classifier to predict PPI is a process that involves using this straightforward probabilistic classification technique. The method operates on the “naïve” assumption that each feature of a class, in terms of its presence or absence, is independent of every other feature. This holds true even if these features are interdependent or if their existence relies on other features. The assumption’s lack of complexity earns the method its “naïve” moniker. Fig. 5 shows the procedure of Naïve Bayes.

**Fig. 5:**
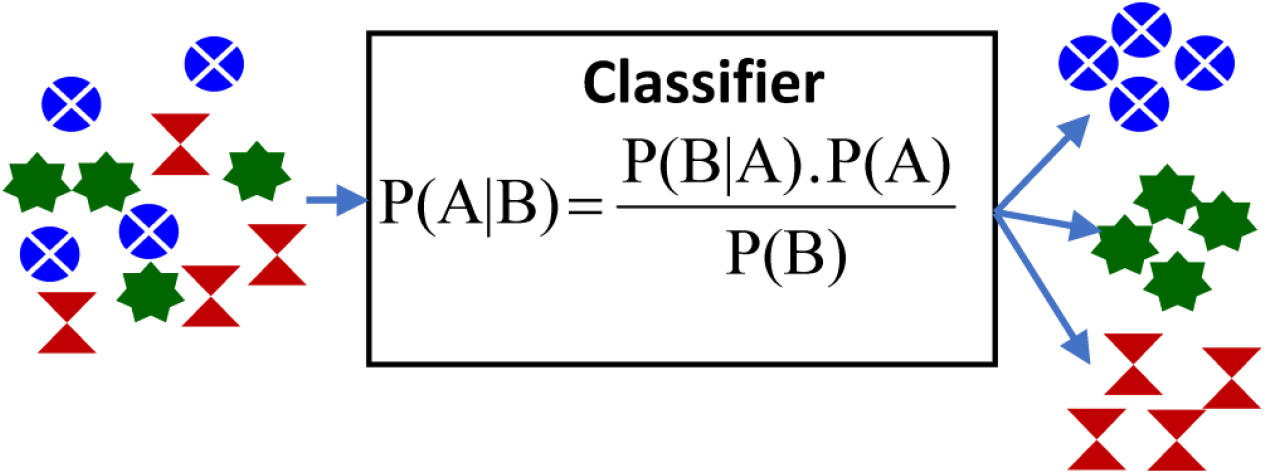
The procedure of Naïve Bayes-based classifier is illustrated in the figure.

Xu et al. [85] addressed the issue of unreliable interactions in protein complexes by employing Fuzzy Naive Bayes, which accounts for unreliability. They concentrated on the classification of protein complexes from subgraphs, taking into consideration the fuzzy characteristic of PPI. Metipatil et al. [86] introduced a collection of machine learning methods, utilizing both Naive Bayes and SVM, to categorize cancer gene information according to sequence-based binary PPI and gene expressions. To enhance different performance aspects like classification accuracy, the authors employed principal component analysis to diminish the dimensionality of the data within these ensemble classifiers.

Hu et al. [87] introduced a strategy that relies on a machine learning algorithm for predicting PPI hotspots. This method involved the training of variables such as amino acid composition, surface area, and amino acid chains. Among various classifiers, the authors incorporated the Gaussian Naive Bayes algorithm into a separate pipeline specifically to forecast protein hotspots. The final outcomes were then refined by employing a combination of voting and stacking schemes.

### 3.3 Instance-Based Model Sub-Category

#### 3.3.1 K-Nearest Neighbor Technique

> Utilizing the K-Nearest Neighbor (KNN) algorithm to predict PPI is an approach that leverages the idea of similarity between features to make predictions. In this method, the feature vectors of proteins, which may include aspects like amino acid sequences, structural properties, and functional domains, are compared. A given protein is then classified according to the majority class of its K nearest neighbors in the feature space, where “nearest” is often measured using Euclidean distance or other distance metrics. The procedure of K-Nearest Neighbor-based classifier is illustrated in Fig. 6.

**Fig. 6:**
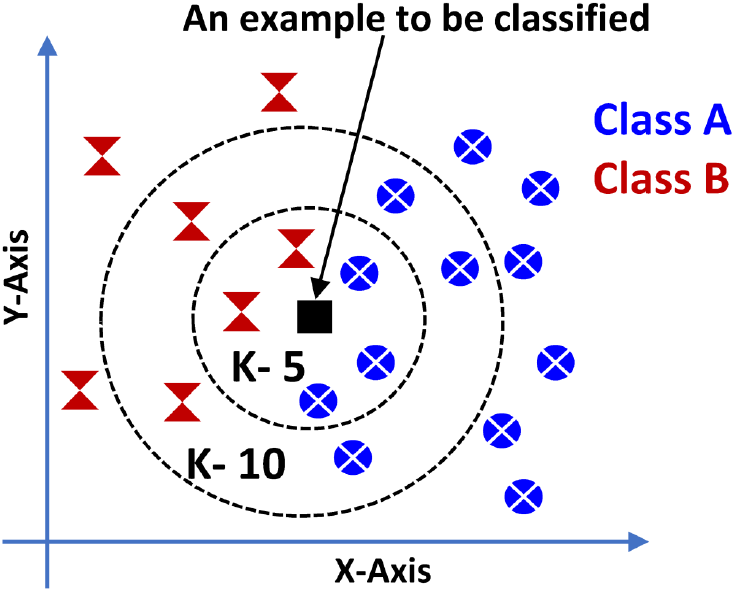
The procedure of K-Nearest Neighbors-based classifier is illustrated in the figure.

Dey and Mukhopadhyay [88] forecasted PPI between dengue and its human host by merging amino acid composition and the conjoint triad of human protein sequences into a feature vector. For prediction, the authors focused on employing techniques such as K-nearest neighbor (KNN), SVM) and Naive Bayes. Shiguihara-Juárez et al. [89] introduced a universal feature representation designed for the PPI task. Initially, parse trees are obtained and labeled with their POS tags. Next, a binary bag-of-words model is formed from these features. Finally, these features are suitable for use with any conventional machine learning method. To test the POS-tag features, the authors employed Weka, utilizing the K-Nearest Neighbor (KNN) classifier, a method solely dependent on distance computation.

Ambert and Cohen [91] showed the effectiveness of a straightforward machine learning method for addressing the issue of identifying PPI. They utilized a version of the kNN classification algorithm in their approach. When selecting the value of k, the algorithm refines this value according to the properties of the training data. It employs a leave-one-out cross-validation cycle on the training data to do so. The value of k is progressively increased by the algorithms, starting from one and continuing up to the total number of samples in the k-training partition of the full data set used for the training of the classifier.

### 3.4 Margin-Based Model Sub-Category

#### 3.4.1 Support Vector Machine (SVM) Technique

> Utilizing Support Vector Machine (SVM) to predict PPI involves using this machine learning model to analyze and understand complex relationships between proteins. In this method, information about proteins, such as sequences and structural features, is converted into feature vectors. These are then used to train the SVM to classify interactions between protein pairs by finding the optimal separating hyperplane. The nonlinear nature of biological data is addressed through various kernel functions, allowing for complex modeling of relationships between proteins. Fig. 7 shows the procedure of Support Vector Machine-based classifier.

**Fig. 7:**
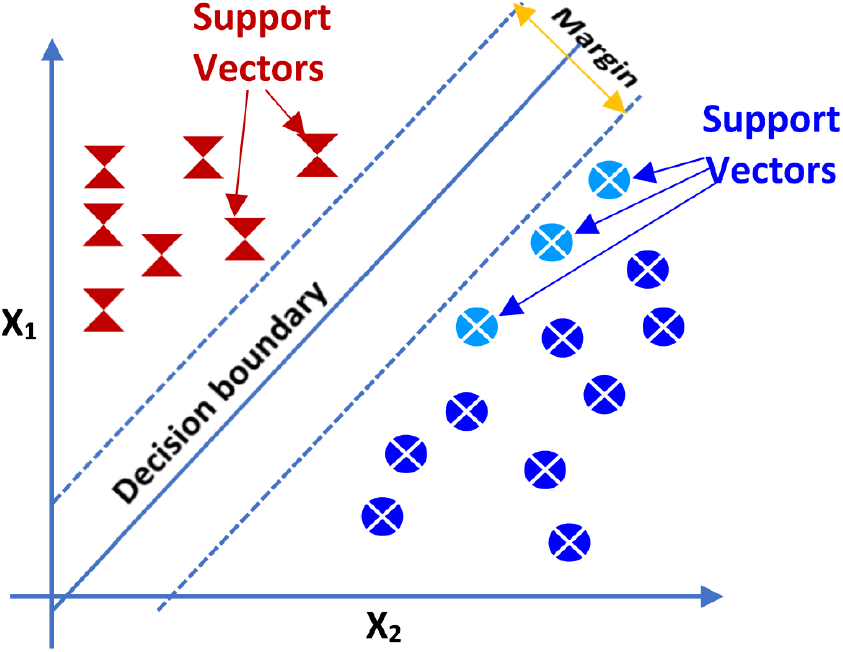
The procedure of Support Vector Machine is illustrated in the figure.

Chen et al. [92] introduced a technique for analyzing biological literature and automatically extracting PPI)relations from texts. This method consists of three main subsystems, organized in a pipeline structure. These subsystems are crafted using machine learning techniques, specifically support vector machines (SVMs) and conditional random fields (CRFs), and are supplemented with thoughtfully constructed informative features. Sætre et al. [93] investigated the variations and commonalities among different PPI approaches. The authors elucidated how to utilize the openly accessible open-source system known as AkaneRE, subsequently detailing the fundamental methods from a scientific standpoint. The AkaneRE system is divided into three components: a core engine for Relation Extraction (RE), a collection of modules designed for solutions, and a configuration language. The core engine employs Support Vector Machines (SVM) and extracts features from the provided training data.

Lin and Zhang [94] introduced a SVM system that relies on ensemble learning for the prediction of hot spot residues. They applied this method to forecast hot regions in PPI. H. Chen et al. [95] introduced a method that utilizes GPU acceleration for SVM, specifically targeting hyperparameter estimation in real-world classification assignments. This approach allows for the efficient and precise training of SVMs, with the computation of kernel functions sped up across various PPI datasets. Arango-Rodriguez et al. [96] suggested using SVM to predict PPI by utilizing physical-chemical characteristics derived from the AA index. Chatrabgoun et al. [97] introduced an algorithm to predict PPI, which estimates the kernel-based SVM by utilizing a low-rank truncated Mercer series. This approach makes the QOP for the approximated kernel-based SVM much more manageable, as there is a substantial decrease in the computational time required for both the training and validation phases.

Karan et al. [98] created a machine learning model utilizing SVM to forecast PPIs between rice and rice blast. The model specifically employs SVM for the detection of these PPIs. Sunggawa et al. [99] utilized SVM as the classification method, and they paired it with four kinds of multivariate mutual information (MMI) to describe amino acid sequences. This approach was used to forecast PPIs, specifically focusing on the interaction data between HIV1 and Humans. The authors based their analysis on the data relating to protein interactions between HIV1 and Humans, which is available in the National Center for Biotechnology Information (NCBI). Li et al. [100] introduced a system for extracting PPIs that relies on the ensemble kernel model combined with active learning. In the initial step, within the SVM classifier, the ensemble kernel is formed by merging the lexical feature-based kernel with the path-based kernel.

Chu et al. [101] implemented a symmetrical pair of feature vectors to illustrate the symmetrical relationship between two proteins. They utilized Sfold cross-validation to identify the optimal SVM parameters. Once determined, these best SVM parameters were then applied to train the SVM-based predictor for PPIs. Feng et al. [102] combined multiple databases containing protein information to predict PPI and build the interaction network. This method was developed using several protein databases and employed a learning algorithm known as SVM.

## 5. Comparative Evaluations

In this section, we examine the various techniques described in this paper, evaluating each one on four key aspects: the underlying principle of the technique, the logic guiding its implementation, the essential conditions needed to reach peak performance, and any constraints. Table 1 delineates the techniques that utilize deep learning for identifying PPI, whereas Table 2 highlights those techniques that employ traditional machine learning for PPI detection. Our objective is to furnish a thorough grasp of the positive and negative attributes of each technique, and to ascertain how suitable they are for particular tasks.

**Table 1.** Evaluating each *<u>deep learning</u>* technique for identifying PPI in terms of the following four criteria: its underlying principle, its justification, its conditions for optimal performance, and its limitations.

| Tech-Papers | Technique<br>Essential Concept | Rationale Behind the<br>Usage of the<br>Technique | Conditions for the<br>Optimal Performance of<br>the Technique | Limitations of the<br>Technique |
| --- | --- | --- | --- | --- |
| Artificial Neural Network (ANN)<br>[26-28] | The method involves creating a network composed of layers containing interconnected nodes or "neurons" that process information related to protein characteristics. The input layer takes various attributes, such as amino acid sequences and structural information, and passes them through hidden layers. During this process, weights and biases are applied, and activation functions transform the data. Through techniques like backpropagation and optimization algorithms, the network is trained on known PPIs. The final output layer provides predictions concerning the likelihood of specific PPIs. | (1) ANNs, known for modeling complex, non-linear functions, making them suitable for PPIs, (2) ANNs' ability to process various protein features (e.g., sequence and structure data) to identify relevant attributes for PPI prediction, (3) ANNs' multi-tiered data representation capability, useful in understanding protein interactions with different abstraction levels, (4) Integration of diverse data types such as genomic information and interaction networks into a unified ANN model for a holistic understanding of PPIs, and (5) Fine-tuning pre-trained models on related tasks for specific PPI prediction, a valuable approach when there's limited labeled data | The enhancement of the method can be achieved through the following: (1) Determining the optimal number of hidden layers and the number of units within each layer, (2) Utilizing suitable activation functions, such as ReLU or tanh, to introduce non-linearity, (3) Using techniques like dropout or L1/L2 regularization to reduce overfitting, (4) Employing fitting algorithms such as Adam or RMSprop, (5) Selecting the best learning rate to ensure steady convergence, (6) Opting for the appropriate batch size and training epochs, and (7) Choosing a relevant loss function, like binary cross-entropy, that is congruent with the problem at hand. | Limitations: (1) Limited validated experimental data leading to overfitting, (2) Need for proper feature selection with deep domain knowledge as incorrect choices can hurt performance, (3) Handling high-dimensional and complex protein data can challenge performance, (4) High computational resources required for training deep ANNs, (5) Sensitivity to noise and errors in experimental data can hinder training and predictions, (6) Lack of generalization for ANNs trained on specific proteins or interactions, limiting broader applicability, (7) Difficulty in integrating various complex data sources, and (8) Time-consuming hyperparameter tuning. |
| Deep Neural Networks (DNN)<br>[29-32] | DNNs consist of several interconnected "neuron" layers, utilizing nonlinear operations to understand PPIs. The network takes varied protein-related data, including sequences, structures, or annotations. Training a DNN on known PPIs involves adjusting the neuron connections through backpropagation, a process that minimizes the error between predicted and actual outcomes, commonly using gradient descent or its variations to iteratively update the model's parameters. Through ongoing refinement, the network can forecast unseen data and potentially uncover new protein interactions. The hidden layers reveal complex features. | DNNs can detect intricate patterns in data, with their ability to discern complexity increasing with network depth. They can engineer features from raw data, identifying nuances. In predicting PPI, DNNs synthesize various information like amino acid sequences, 3D structures, and functional annotations, enhancing prediction accuracy. Different DNN architectures can be used for specific tasks in PPI prediction. This includes analyzing sequences, handling sequential data, and considering interconnected protein interactions. DNNs can be adapted for specialized tasks, reducing training time and leveraging prior knowledge. Their global optimization ability is essential for understanding complex protein relationships. | The method can be enhanced by: (1) Balancing simplicity and complexity in modeling to capture PPIs without causing overfitting. (2) Carefully selecting and possibly tuning Batch Size and Learning Rate through cross-validation. (3) Using L1 or L2 regularization, or dropout, to prevent overfitting. (4) Managing model complexity to evade both underfitting and overfitting. (5) Selecting the right network architecture, like CNNs or RNNs. (6) Incorporating known biological relationships and constraints. (7) Applying early stopping or utilizing validation sets in training. (8) Using appropriate evaluation metrics tailored to the data imbalance. (9) Implementing cross-validation or other validation strategies. | Limitations: (1) The need for large, labeled datasets that are costly and time-consuming to generate and may lack coverage; (2) The risk of learning biases from uneven data representation, causing inaccuracies; (3) The complexity of handling high-dimensional protein data; (4) Significant computational resource requirements, including specialized hardware; (5) Risk of overfitting to noise due to numerous parameters; (6) Difficulty in integrating diverse influencing factors like genetic and environmental aspects; (7) Optimization challenges in training, such as getting stuck in local minima; (8) Ensuring that the model generalizes across different biological contexts is a challenge. |
| Extreme Learning Machine (ELM)<br>[33-40] | ELM is unique in its approach to predicting PPI by randomly initializing the weights and biases in the hidden layer, then determining the output weights analytically instead of through iterative adjustments. This accelerates the training process, making it suitable for handling complex datasets related to PPI. ELM's one-step learning process eliminates the iterative tuning, achieving quicker learning without a reduction in performance, as it has been found to deliver equal or superior prediction accuracy compared to other techniques. | ELM is celebrated for its rapid learning and simple design, utilizing randomly assigned hidden weights, which aids in handling vast PPI data. Its single hidden layer models complex non-linear relationships in PPIs, while its structure helps resist overfitting in situations where parameters exceed data points. ELM's adaptability allows it to handle diverse data essential for predicting intricate PPI. Demonstrating robust generalization, it discerns underlying patterns without extensive tuning. ELM can be combined with other models for better accuracy, and customized to manage imbalanced data. | The method can be enhanced by: (1) Adjusting input features to fall within a similar range, (2) Choosing the right kernel function, such as RBF or linear, (3) Properly determining the number of hidden neurons, incorrect amounts can hinder performance, (4) Considering the activation function, (5) Designing the ELM architecture to match specific needs of PPI data, (6) Using Early Stopping to prevent overfitting by stopping training when no further improvement on a validation set, (7) Merging multiple ELM models to capture various aspects of data, (8) Applying optimization techniques to shorten training without sacrificing accuracy. | ELM's one hidden layer may hinder complex pattern detection in PPI. Its static weight initialization between input and hidden layers may cause inconsistency and suboptimal results. The absence of a backpropagation phase limits adaptiveness, potentially affecting intricate tasks like PPI prediction. ELM may overfit on small or complex datasets, and it's challenged by the imbalance in PPI data. Finally, its performance can be sensitive to hyperparameter adjustments, such as hidden node count, activation functions, and regularization, where minor changes may lead to significant outcome variations. |
| **Convolutional Neural Network**<br>[41-44] | The technique uses a filter or kernel to slide over an input matrix (e.g., protein sequence) to form a feature map, preserving pixel relationships by detecting image attributes with small input data portions. After each convolution, a nonlinear layer like ReLU introduces nonlinear aspects. Pooling reduces feature map size but retains essential information. After convolution and pooling, the design features fully connected layers where the matrix connects to neurons in a dense layer, enabling the network to interpret complex input features and make predictions | Proteins interact through complex 3D structures, and CNNs are designed to handle these spatial relationships. CNNs use multiple layers to capture varying levels of abstraction, with lower layers identifying basic structural motifs and higher layers discerning complex protein interactions. Some CNN architectures allow multi-scale representation to recognize local and global patterns, essential for understanding PPI. CNNs can integrate various information such as sequence data and structural details, enabling accurate modeling of protein interactions. They also balance fitting training data with generalizing to new examples. | The method can be improved by: (1) Encoding proteins with one-hot encoding or embedding layers to convey sequence details, (2) Enhancing robustness through transformations like rotation and flipping, (3) Properly designing convolutional, pooling, and fully connected layers, (4) Selecting appropriate filter sizes for spatial patterns, (5) Using functions like ReLU for non-linearity, (6) Applying a suitable loss function like binary cross-entropy, (7) Using techniques like dropout and L2 regularization to prevent overfitting, and (8) Applying attention to focus on key sequence areas. | Limitations: (1) CNNs, designed for 2D data, face challenges in accurately representing complex 3D protein structures, causing difficulties in processing protein interactions, (2) Recognizing long-range dependencies between different parts of a protein can be problematic for CNNs, as convolutional layers may miss key global and spatial interactions, (3) it requires substantial labeled data, (4) Training complex CNN models for PPI prediction is computationally demanding, (5) Proteins' diversity means a model trained on specific proteins may not generalize well, becoming too specialized to the training data |
| **Graph Convolutional Networks (GCN)**<br>[45-48] | GCNs carry out convolution by collecting data from adjacent nodes, using local amalgamation to merge information from the neighbors of a node. This method can encapsulate the local arrangement around each protein. The procedure of graph convolution broadens the conventional convolution concept, moving it from grid-based data to graph structures. It modifies the features at a specific node by gathering attributes from its neighboring nodes, symbolizing the mutual influence between proteins. Comprising several layers of graph convolution, it captures various degrees of abstraction within the graph. | GCNs are suited for predicting PPIs, as they can represent proteins as graphs where nodes are amino acid residues, and edges symbolize spatial relationships. These interactions between proteins can be modeled as graph interactions. Designed to work with these structures, GCNs consider both local and global structural details, understanding the spatial relationships within a protein and interaction patterns between proteins. This enables an effective prediction of PPIs, capturing the full complexity of the molecules. By adapting the convolution operation for graphs, GCNs learn essential protein features from their graph representation. | The method can be enhanced by: (1) Careful choice of the quantity and variety of convolutional layers, (2) Implementation of suitable activation functions and non-linearities, (3) Employment of regularization methods such as dropout to avoid overfitting, (4) Selection of a loss function that is specifically adapted to the task, (5) Utilization of an optimal optimization algorithm, fine-tuned with the right hyperparameters, (6) Integration of ensemble or hybrid models that merge GCNs with different architectures, (7) If relevant, the application of transfer learning, (8) Efficient management of unbalanced data and the enhancement of data through augmentation. | Limitations: (1) GCNs might not fully understand the complex factors of protein interactions, especially when information is lacking, (2) They can be slow and computationally heavy, particularly with large interaction networks, (3) Accurate modeling requires abundant quality data; poor data may cause imprecise predictions, (4) Models specializing in specific proteins or interactions may not adapt well to new types due to proteins' varied functions and structures, (5) GCNs may be sensitive to noise, resulting in wrong predictions, (6) Integrating various data like sequence and structural information into GCNs might be complex. |
| **Autoencoder**<br>[49-56] | An autoencoder works by compressing high-dimensional features into a lower-dimensional latent space through an encoding process, using a series of nonlinear transformations that are learned from the data. The encoded form captures the essential information needed to represent the proteins. The decoding part works to reconstruct the original high-dimensional data from this compressed representation. By training to minimize the difference between the original data and its reconstruction, the network learns to capture the most important aspects of the data in the compressed form | (1) Autoencoders condense complex protein information to simpler form, keeping essential features, (2) They interpret protein interactions, such as amino acid sequences and structures, (3) Being unsupervised, they don't need labeled data, suitable for analyzing large amounts of biological data, (4) They handle various data types, combining them into cohesive representation, (5) As neural networks, they model complex relationships between variables, (6) They can be fine-tuned to emphasize critical data features, (7) The encoder part can be adjusted for targeted predictions, building on properties of proteins learned unsupportively | Enhance the method by: (1) Identifying the optimal number of hidden layers and units to balance data complexity and overfitting; (2) Using L1 or L2 regularization, dropout layers, etc., to avoid overfitting; (3) Selecting the right batch size for training speed and convergence stability; (4) Determining the proper number of epochs to avoid underfitting or overfitting; (5) Applying advanced optimization algorithms like Adam, RMSProp for faster convergence; (6) Integrating autoencoders with models like SVM to enhance performance; (7) Systematically searching hyperparameter space using techniques like Grid Search or Random Search for improvement. | Limitations: (1) Autoencoders reduce data into a lower-dimensional space and then reconstruct it, potentially losing vital information for predicting PPIs, (2) Mainly used for encoding and decoding data, autoencoders don't have mechanisms to model protein interactions, key in PPI prediction, (3) Training autoencoders can be time-consuming with large datasets, (4) The aim of minimizing the difference between input and output in autoencoders doesn't align with predicting PPI, (5) Depending on training, an autoencoder may not adapt well to unseen proteins and interactions, only reflecting training data characteristics. |
| **Long Short-Term Memory (LSTM)**<br>[63-67] | LSTMs incorporate a unique structure of gates and memory cells that allow them to effectively learn and remember relevant information across long sequences. LSTM utilizes a forget gate to decide what information to erase from its memory, an input gate to update its memory, and an output gate to control what information is passed on to subsequent layers. LSTM's ability to remember intricate sequence patterns and long-range dependencies enables it to model complex relationships between amino acids, which in turn can reveal how proteins might interact with each other | LSTM are ideally suited for handling sequential data like protein sequences; they can remember and utilize long-range dependencies within those sequences for accurate predictions; they aid in understanding the 3D structure of proteins, crucial for PPI; they can simultaneously handle multiple related tasks, enhancing prediction accuracy; they generalize well on unseen data, making them useful for new proteins; and they can integrate various features such as physicochemical properties and known pathways for a comprehensive understanding of PPI. | Improving the method can be done by: (1) Creating a design with the right LSTM layers, units, and dropouts, and using bi-directional LSTMs for sequence information, (2) Adding an attention mechanism for network focus on key sequence parts, (3) Selecting an apt loss function, (4) Utilizing dropout or L2 regularization to avoid overfitting, (5) Adjusting the learning rate during training for faster convergence, (6) Monitoring validation loss to stop training at performance plateaus, avoiding overfitting, and (7) Merging LSTM with architectures like CNNs or Graph Neural Networks to capture more data patterns. | Limitations: (1) Training and fine-tuning LSTM networks is challenging due to their complex gating mechanisms. This can lead to higher costs and longer training times, (2) LSTMs may not efficiently capture spatial relationships within proteins, (3) LSTMs may suffer from the vanishing gradient problem with long sequences, (4) LSTMs may not fully grasp 3D spatial relationships, vital for accurate PPI prediction, (5) LSTMs' flexibility can lead to overfitting if not properly regularized, especially with small training datasets, (6) Generalizing LSTMs across different protein types might require re-training or fine-tuning. |

**Table 2.** Evaluating each *<u>traditional machine learning</u>* technique for identifying PPI in terms of the following four criteria: its underlying principle, its justification, its conditions for optimal performance, and its limitations.

| **Tech.** | **Papers** | **Technique Essential Concept** | **Rationale Behind the Usage of the Technique** | **Conditions for the Optimal Performance of the Technique** | **Limitations of the Technique** |
| --- | --- | --- | --- | --- | --- |
| **eXtreme Gradient Boosting (XGBoost)** | **[68-73]** | XGBoost constructs a sequence of decision trees, with every new tree aiming to rectify the errors of the combined set of existing trees. This lessens a loss function. Hyperparameters like the learning rate, the depth of the trees, and the regularization elements, can be adjusted to match the unique characteristics of PPI data. The model employs a depth-first strategy and trims the trees using the 'max_depth' attribute. This facilitates increased complexity in the necessary areas without unduly extending other segments of the tree. | (1) XGBoost is suitable for PI prediction due to its scalability and efficiency in handling large datasets, (2) As an ensemble method, it leverages multiple decision trees to uncover complex patterns in PPIs, (3) It employs L1 and L2 regularization to prevent overfitting, improving generalization, (4) Its ability to handle missing data adds flexibility to the training process, (5) Its feature-ranking capability helps researchers uncover what influences predictions, (6) The tunable hyperparameters in XGBoost facilitate precise model adjustment, crucial for PPI prediction. | The method can be enhanced by: (1) Tuning parameters such as the learning rate, tree's maximum depth, number of estimators, and subsample ratio to suit the particular dataset, (2) Employing suitable regularization methods like L1 or L2 to prevent overfitting, (3) Introducing early stopping to mitigate overfitting by observing a chosen metric on a dataset, (4) Utilizing the capability of XGBoost to run on multiple CPUs and GPUs, which can cut down training times, (5) Pairing XGBoost with other machine learning models for PPI prediction can further improve performance. | Limitations: (1) Struggling with high-dimensional data, leading to overfitting, especially with limited data, (2) Sensitivity to imbalance in PPI datasets, causing skewed predictions, (3) High computational expense, (4) A demanding process for tuning numerous hyperparameters, (5) Potential overlooking of intricate interdependencies in proteins, leading to decreased accuracy, (6) Lack of consideration for sequential dependencies in protein structures, (7) Incompatibility with the natural integration of varied biological data types like genomic or proteomic information. |
| **Gradient Boosting (G-Boosting)** | **[74-76]** | The boosting process starts by fitting a simple model to the data and calculating the prediction error for each observation. Then, a new decision tree is fit to these errors, trying to predict how much each observation's prediction should be adjusted. The predictions are scaled by a learning rate, and then added to the original predictions. This process is repeated, each time fitting a new tree to the residual errors of the combined ensemble of trees, until the improvement in prediction plateaus. | G-Boosting can avoid overfitting by adjusting hyperparameters like learning rate and tree depth. In PPI, often having non-linear relationships, gradient boosting models these without direct definition, improving predictive efficiency. A challenge in PPI data is handling missing data; G-Boosting does this without imputation, avoiding bias. Its flexibility in using various loss functions tailors to PPI prediction needs. G-Boosting's ability to merge diverse data such as sequence details and functional annotations is essential in PPI prediction. | The method can be enhanced by: (1) Choosing a loss function that aligns with the nature of predicting PPI, such as using binary cross-entropy for binary classification, (2) Carefully determining the optimal number and depth of trees in the ensemble to balance complexity with the risk of overfitting, (3) Modifying the learning rate to regulate the influence of each individual tree, (4) Utilizing hyperparameter tuning techniques like Random Search, (5) For training on extensive datasets, considering approaches for parallelization. | The method faces challenges due to its complexity, such as difficulty in interpretation and potential overfitting without precise tuning. Training can be computationally demanding, especially with large, multidimensional data. Issues may arise with noisy PPI data, sensitivity to outliers, and uncertainty about the importance of specific features. It may not perform well on imbalanced data, and predicting PPI may be complicated due to the diverse data involved. Capturing non-linear relationships requires careful hyperparameter tuning, and optimizing it is time-consuming. |
| **LightGBM** | **[24, 77, 78]** | LightGBM operates by constructing a series of decision trees, where each tree corrects the errors of its predecessors, iteratively improving the model's predictions. The gradient-based one-side sampling (GOSS) and exclusive feature bundling (EFB) techniques within LightGBM reduce the data and feature redundancy, further improving the efficiency. GOSS keeps the instances with large gradients and randomly samples the instances with small gradients, while EFB bundles exclusive features together to reduce the dimensions. | (1) It uses histogram-based algorithms to speed up training and reduce memory use; (2) It achieves similar accuracy to other algorithms but with less computing resources, important for complex PPI; (3) It can handle categorical features without preprocessing; (4) It offers customization for PPI prediction, including custom functions and criteria; (5) It supports multi-class classification, suitable for various interaction types in PPI; (6) It includes L1 and L2 regularization to prevent overfitting; (7) It can be easily integrated into existing bioinformatics workflows, adding to its adaptability. | The method can be improved by: (1) selecting the right Learning Rate, with smaller rates potentially requiring more iterations; (2) using techniques like SMOTE for class imbalance in PPI prediction; (3) controlling tree depth to prevent overfitting; (4) setting a minimum data limit in a leaf to avoid overfitting; (5) choosing an appropriate boosting strategy, such as 'gbdt'; (6) letting LightGBM handle categorical features without one-hot encoding; (7) aligning the objective function and metric with PPI prediction; (8) automating hyperparameter tuning; (9) using stacking or bagging with LightGBM | Limitations: (1) LightGBM can handle some non-linearity, but may be challenged by complex biological interactions, (2) If overly intricate and with a small dataset, LightGBM may overfit the training data, (3) LightGBM is sensitive to noise, (4) Despite its efficiency compared to other models, LightGBM can be resource-intensive, especially with large datasets or hyperparameter tuning, (5) LightGBM might struggle with imbalanced data without proper handling, (6) Its inability to integrate diverse data types may limit LightGBM's effectiveness in PPI prediction |
| **Random Forest** | **[79-82]** | It starts by selecting random samples from the training dataset. On each of the dataset subsets, a decision tree is built. At each node, only a random subset of the features is considered for splitting. The method combines multiple decision trees, each built on a random subset of the features, to form a "forest." Each tree in this forest gives a classification, and the mode of the classes from individual trees is taken as the final prediction. The individual trees "vote" on a prediction, and the majority class is taken as the final prediction | (1) Utilizing a voting system among the trees, the final prediction is enhanced in accuracy and robustness compared to depending on a single model, (2) Random Forest's efficiency in managing high-dimensional data eliminates the need for intricate feature selection, (3) Training each tree with a unique data subset reduces overfitting, (4) Random Forest identifies nonlinearities without assuming specific data distribution or model function, (5) The ensemble approach makes Random Forest relatively resistant to noise, helping mitigate individual data discrepancies. | Refining a Random Forest necessitates confronting issues such as unequal datasets, picking relevant attributes, data normalization, and dealing with incomplete values. Adjusting essential hyperparameters like the count of trees, the depth of the trees, and the criteria for splitting is crucial to avoid overfitting and sharpen the model's efficiency. Attention must also be given to computational factors and parallel computation might be utilized. The model's efficacy may be further augmented by combining it with other algorithms or employing methods of regularization. | Limitations: (1) Struggling with irrelevant or missing features, often requiring domain knowledge to select proper ones; (2) Consuming high computational resources with large datasets, leading to increased training time; (3) Overfitting if not tuned correctly in complex PPI; (4) Risk of bias towards the majority class in imbalanced datasets, affecting prediction for the minority class; (5) Difficulties in handling heterogeneous data types without significant pre-processing; (7) Sensitivity to hyperparameter choices, with the tuning process being time-consuming and requiring expertise. |
| **Probabilistic Decision Tree (PDT)** | **[25, 83, 84]** | A typical PDT is built by analyzing known PPI and using statistical measures to estimate the likelihood that two proteins will interact based on various features, such as their physical properties, biological functions, and structural characteristics. The tree's structure is defined by decision nodes, where each decision is based on a specific feature or set of features that may influence the interaction. As one traverses the tree, the decisions and corresponding probabilities lead to a final prediction regarding the probability of interaction. | Decision trees' ability to process probability captures the complexity in PPI prediction, allowing for nuanced modeling. PDTs simplify the examination of interactions and facilitate the integration of diverse data like sequence and structural information without complication. They assess various likely hypotheses, making the model resilient to data's noise. PDTs provide a visual representation of decision-making, boosting researchers' understanding into biological phenomena. Their ability to seamlessly incorporate different data and adapt to various contexts makes them a versatile tool tailored to various research needs. | The choice of relevant features is critical to building a model that is neither too complex nor too simple. Including features that contribute to PPI without adding redundant or irrelevant information ensures better performance. Model hyperparameters should be carefully tuned. This involves choosing the correct structure of the tree, including the depth, splitting criteria, and leaf node thresholds. Handling imbalanced data is essential if there are disproportionate numbers of interacting versus non-interacting protein pairs. Techniques to balance the data or adjust the algorithm's sensitivity may be required. | Limitations: (1) PDTs can't capture the nonlinear, context-specific nature of PPI influenced by various factors, reducing them to binary decisions with set rules, (2) The independence assumption in PDT may ignore complex dependencies between features in data, (3) The accuracy of PDT depends on the quality and quantity of the training data, and issues like overfitting may arise from noisy data, (4) The probabilistic aspect of PDT can introduce uncertainties, leading to incorrect conclusions if not correctly estimated, (5) PDT may not be suitable for large-scale PPI prediction, as it can be computationally expensive |
| **Naïve Bayes** | **[85-87]** | The model computes the probability of interaction between two proteins based on the observed features. It calculates the conditional probability of the interaction given the observed features. The probabilities are estimated from a training set of known PPI. The model will then classify a new pair of proteins as interacting or not interacting based on which probability is higher. It does this by comparing the conditional probabilities of the two classes, given the observed features for that particular pair of proteins. | The rationale for using Naïve Bayes in predicting PPI comes from its probabilistic approach and computational efficiency. Its assumption that all features are independent given the class variable makes it suitable for high-dimensional feature spaces found in protein studies. This independence, although simplistic, helps avoid the curse of dimensionality, making the method scalable. Naïve Bayes' probabilistic nature offers insights by giving confidence scores to predictions, aiding in decision-making. It can also handle missing data, common in biological datasets. | The optimal performance requires careful consideration of several factors. The feature selection must be relevant to represent the proteins, and the model's assumption that these features are independent should be met. Choosing the right probability distribution, such as Gaussian, Multinomial, or Bernoulli, and proper preprocessing like normalization and scaling may be necessary based on the data's nature. Hyperparameter tuning through methods like cross-validation helps find the best configuration, and the integration of biological context can lead to more accurate predictions of PPI. | Limitations: (1) The fundamental assumption that features are independent often doesn't hold with complex protein interactions, leading to prediction inaccuracies, (2) Its simple probabilistic nature may result in the inability to capture complex, non-linear relationships, leading to oversimplification and reduced predictive accuracy, (3) Including unrelated information can distort predictions, (4) The model's effectiveness relies heavily on balanced, representative, and quality training data; imbalanced, biased, or noisy data can make predictions misleading. |
| **K- Nearest Neighbor** | **[88-91]** | The method classifies a protein interaction based on the interactions of its K most similar or 'nearest' neighbors in the feature space. These neighbors are identified based on a defined distance metric, such as Euclidean distance, that quantifies the similarity between proteins. The algorithm looks for the K closest known interactions in this multidimensional space and makes a prediction based on the majority class of these neighbors. If, for instance, most of the K nearest known interactions are positive, then the algorithm predicts a positive interaction for the unknown pair | KNN is an instance-based learning method that classifies new instances by the majority class of the 'K' nearest neighbors from the training data. This approach is beneficial for complex data like PPIs, where the governing function may be unknown. KNN works in a multi-dimensional space representing protein features, capturing local similarity to identify likely interactions. Its simplicity and computational efficiency enable processing of large datasets common in proteomics, providing a more accurate representation of proteins and potential interactions. | Optimal model performance requires a combination of appropriate parameter tuning and data preprocessing. Selecting the correct value for 'K', or the number of neighbors to consider, is crucial, as it can influence the algorithm's bias-variance trade-off. The choice of distance metric is vital, with options like Euclidean and Manhattan distance often used. Normalizing or scaling the features can be essential, as it ensures that all dimensions contribute equally to distance calculations. Cross-validation can help in fine-tuning the hyperparameters to find the optimal set that delivers the best predictive performance. The model yields good results when combined with other techniques. | The method's effectiveness is greatly influenced by the choice of distance metric and k value, potentially causing incorrect predictions. In high-dimensional spaces, like PPI data, it may suffer from the curse of dimensionality, leading to overfitting and high computational costs. KNN's performance depends on the quality of protein feature representation and doesn't inherently offer confidence scores. Its computational cost can be significant with large datasets, and as a lazy learning method, it can memorize data, leading to poor generalization and vulnerability to noise. |
| **Support Vector Machine (SVM)** | **[92-102]** | SVM is a popular supervised learning model for predicting PPI. SVM works by finding a hyperplane that best divides a dataset into classes, maximizing the margin between the closest points of different classes, known as support vectors. The SVM models the relationships between features and the known interactions using a kernel function, allowing for non-linear relationships to be captured by mapping the data into a higher-dimensional space where it becomes linearly separable. By finding the optimal hyperplane, SVM can generalize well from known interactions to make predictions on unseen data. | SVMs are known for handling high-dimensional data, allowing various features in the model. This leads to accurate predictions. Kernels like linear, polynomial, or RBF transform the feature space, making data more separable and capturing complex relationships between protein features. SVMs resist overfitting, especially with proper regularization. This helps the model learn the underlying structure of protein interactions. The SVM's optimization strategy maximizes the margin between classes, focusing on difficult cases, thereby creating a model sensitive to subtle distinctions between proteins, essential for accurate PPI modeling. | The optimal performance depends on selecting the right kernel functions like linear, polynomial, or RBF, and tuning parameters such as regularization (C). This process aids in data transformation and fine-tuning, preventing overfitting and under-fitting. Feature selection and preprocessing techniques, including normalization and handling missing data, enhance the model by focusing on relevant attributes for PPI prediction. Reducing dimensionality is also essential. Integrating biological information like GO annotations or combining with other machine learning techniques elevates prediction accuracy, cementing SVM as a powerful tool in this area. | SVMs are sensitive to kernel selection, with incorrect choices causing overfitting and poor generalization. They often need extensive labeled data, a challenge for protein-protein interactions (PPI). Performance may suffer with imbalanced or sparse data, and high dimensionality can make them slow to train. They also require data to lie in a metric space, and if this assumption is not met, performance can suffer. For complex, nonlinear PPI prediction, traditional linear SVM may be ineffective, necessitating more advanced techniques. SVMs also lack interpretability, making it difficult to understand why a particular prediction was made. |

## 6. Experimental Evaluations

In this section, experiments are conducted to scrutinize and rank the various methodologies delineated in this paper. Each group of algorithms that employs the same core technique is represented by one selected algorithm. These chosen algorithms were then systematically evaluated and ranked. All the algorithms were executed on a Windows 10 Pro system, equipped with an Intel(R) Core(TM) i7-6820HQ processor running at 2.70 GHz, and supported by 16 GB of RAM.

### 6.1 Datasets

- *Database of Interacting Proteins (DIP)* [86]: This PPI dataset was collected from the core subset of Saccharomyces cerevisiae in the DIP. This dataset encompasses 5221 proteins and 24918 interactions, all gathered from 18229 different experiments. DIP serves as a biological repository that compiles PPI confirmed through experimentation. It integrates data from diverse sources to formulate a unified and consistent collection of protein interactions. The interactions included in DIP are ascertained through various experimental methods, including yeast two-hybrid screening, affinity purification, co-immunoprecipitation, and more.
- *Human Protein Reference Database (HPRD)* [23]: This dataset has been developed to aid in the research of proteins within human biology. It delivers extensive and meticulously curated details on PPIs, along with other relevant information concerning human proteins. It contains 30,047 proteins and 41,327 interactions.
- *Search Tool for the Retrieval of Interacting Genes/Proteins (STRING)* [89]: It is a resource that provides a comprehensive and critical assessment of protein-protein interactions. It includes both physical and functional associations, and the interactions are derived from various sources, including experimental data and computational prediction methods.

### 6.2 Evaluation Setup

#### 6.2.1 Preprocessing

- *Feature Extraction:* Extracted the following features: sequence information, structural features, evolutionary information, one-hot encoding, etc.
- *Feature Scaling:* Applied Min-Max normalization.
- Data Augmentation: Applied rotation and flipping.
- *Data Split:* We divided the datasets as follows: 70% training, 15% validation, and 15% testing.

#### 6.2.2 Network Architecture

- *Number of Layers:* 3.
- *Number of Units in Each Layer*: 100.
- *Activation Functions:*

­ Hidden layers: ReLU and tanh,
­ Output layer: Sigmoid
- *Loss Function:* Binary Cross-Entropy.
- *Regularization Methods:*

­ Dropout: 0.5
­ L2 regularization with λ = 0.0005

#### 6.2.3 Training Parameters

- *Batch Size:* 64.
- *Number of Epochs:* 100.
- *Learning Rate:* 0.001.
- *Optimization Algorithm:* Adam.
- *Regularization*: L2 regularization to prevent overfitting.
- *Early Stopping Criteria:* No improvement in validation loss for 10 consecutive epochs.
- *Loss Function:* Cross-entropy loss for classification of interactions.

#### 6.2.4 Evaluation Metrics

- *Sensitivity:* It measures the proportion of actual interacting protein pairs (true positives) that were correctly identified as such by the model. A higher sensitivity indicates that the model is good at detecting actual interactions and not missing them. The formula for sensitivity is defined below: Sensitivity = True Positives / (True Positives + False Negatives)

­ True Positives (TP): The number of correctly predicted interacting protein pairs.

­ False Negatives (FN): The number of actual interacting protein pairs that were incorrectly predicted as non-interacting

- *Specificity:* It is the proportion of actual non-interacting pairs (true negatives) that are correctly identified as non-interacting by the model. A high specificity means that the model is good at correctly identifying non-interacting pairs, minimizing the risk of false alarms (false positives). This is particularly important in PPI prediction, where falsely predicting an interaction between two proteins can lead to incorrect biological understanding and potentially misguide subsequent experimental work. The formula for specificity is: Specificity = True Negatives / (True Negatives + False Positives)

­ True Negatives (TN): The number of actual non-interacting protein pairs that were correctly identified as non-interacting.

­ False Positives (FP): The number of actual non-interacting protein pairs that were incorrectly identified as interacting.

- Precision: It is the proportion of true positive interactions among all the predicted positive interactions. High precision indicates that a greater proportion of the predicted interactions are actual interactions. A model with high precision is cautious in predicting interactions and tends to make fewer false positive errors. In PPI prediction, precision is particularly important when the cost of false positives is high. For example, if predicting a false interaction might lead to costly and time-consuming experiments to verify it, high precision would be desirable to minimize such unnecessary efforts. The formula for precision is: Precision = True Positives / (True Positives + False Positives)

­ True Positives (TP): The number of correctly predicted interacting protein pairs.

­ False Positives (FP): The number of non-interacting protein pairs that were incorrectly identified as interacting.

#### 6.2.5 Experimental Controls

- Cross-Validation: Performed 5-fold cross-validation.
- Hyperparameter Tuning: Used Grid Search tool to find the best hyperparameters.
- Experiment Repetition: For statistical robustness, we performed the whole experiment 5 times and reported the mean and standard deviation of the results.

### 6.3 Methodology for Selecting a Representative Algorithm for Each Technique

The following approach was employed for conducting the experimental evaluations:

- **Evaluating each sub-technique:** After conducting an in-depth examination of papers detailing algorithms that utilize a particular sub-technique, we were able to identify the paper with the greatest impact. This paper’s algorithm was chosen as the representative of the sub-technique. To determine the most significant paper among all those describing algorithms that use the same sub-technique, we assessed various aspects such as the degree of innovation and the date of publication. Table 3 displays the list of the papers we selected.
- **Ranking the sub-techniques that belong to the same overall technique:** We computed the average scores of the selected algorithms that utilized a particular sub-technique in common. Then, we ranked the sub-techniques, all part of the same larger technique, according to these scores.
- **Ranking the various techniques that belong to the same sub-category:** We calculated the average scores for the selected algorithms using the same technique. Then, we arranged the techniques within the same sub-category by ranking them according to these scores.
- **Ranking the various sub-categories that belong to the same category:** We calculated the average scores for the selected algorithms that made use of the same sub-category. Following this, we arranged the sub-categories within the same category in rank order based on these scores.

**Table 3.**
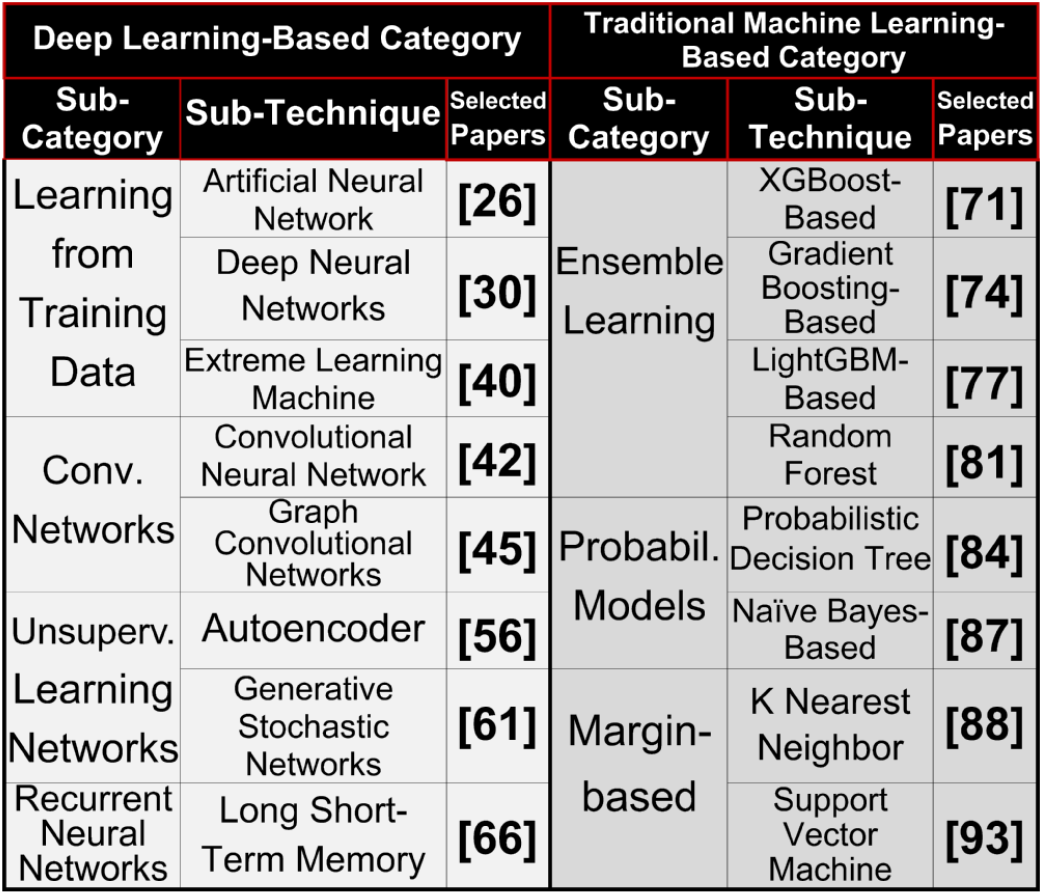
List of selected representative papers.

**Table 4.**
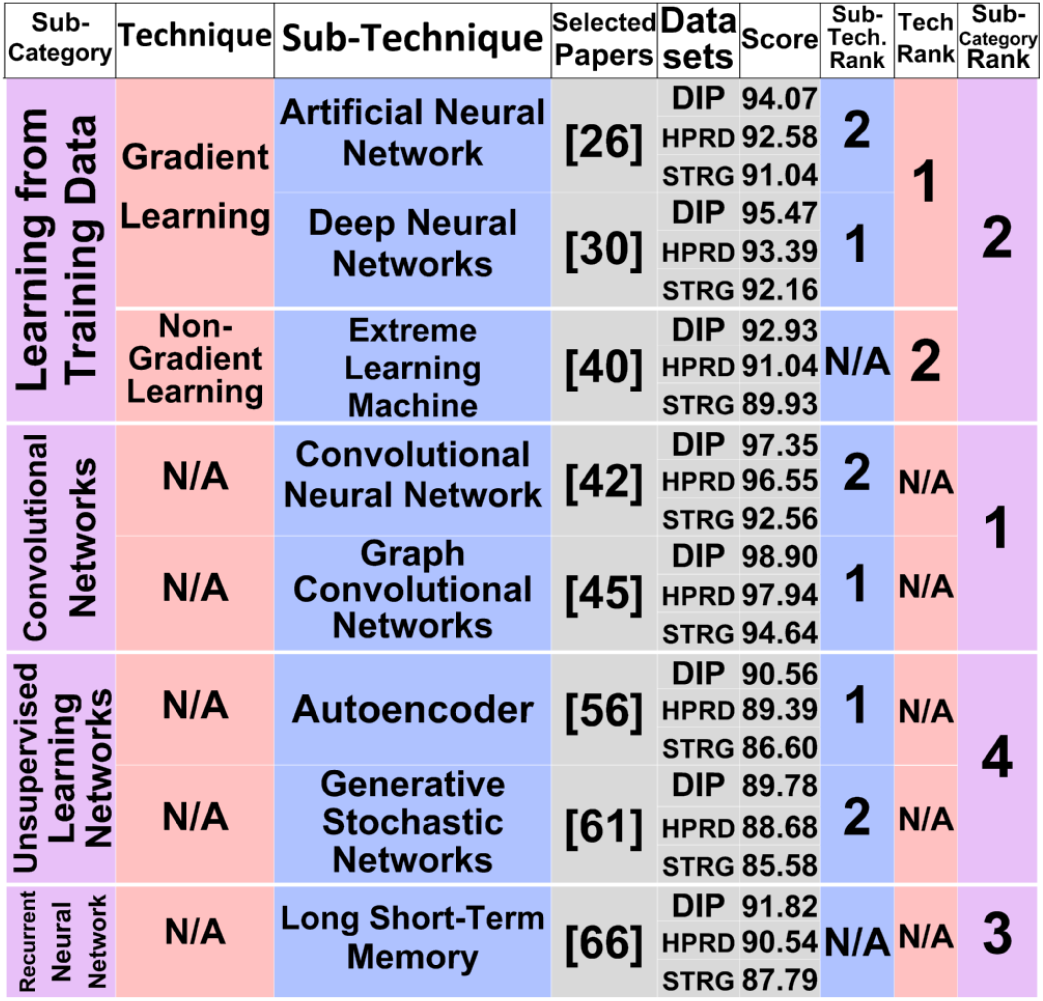
<u>Sensitivity</u> scores of the selected <u>Deep Learning algorithms</u>. The table also shows the following: the ranking of the sub-techniques that belong to the same overall technique, the ranking of the various techniques that belong to the same sub-category, and the ranking of the various sub-categories that belong to the same category.

**Table 5.**
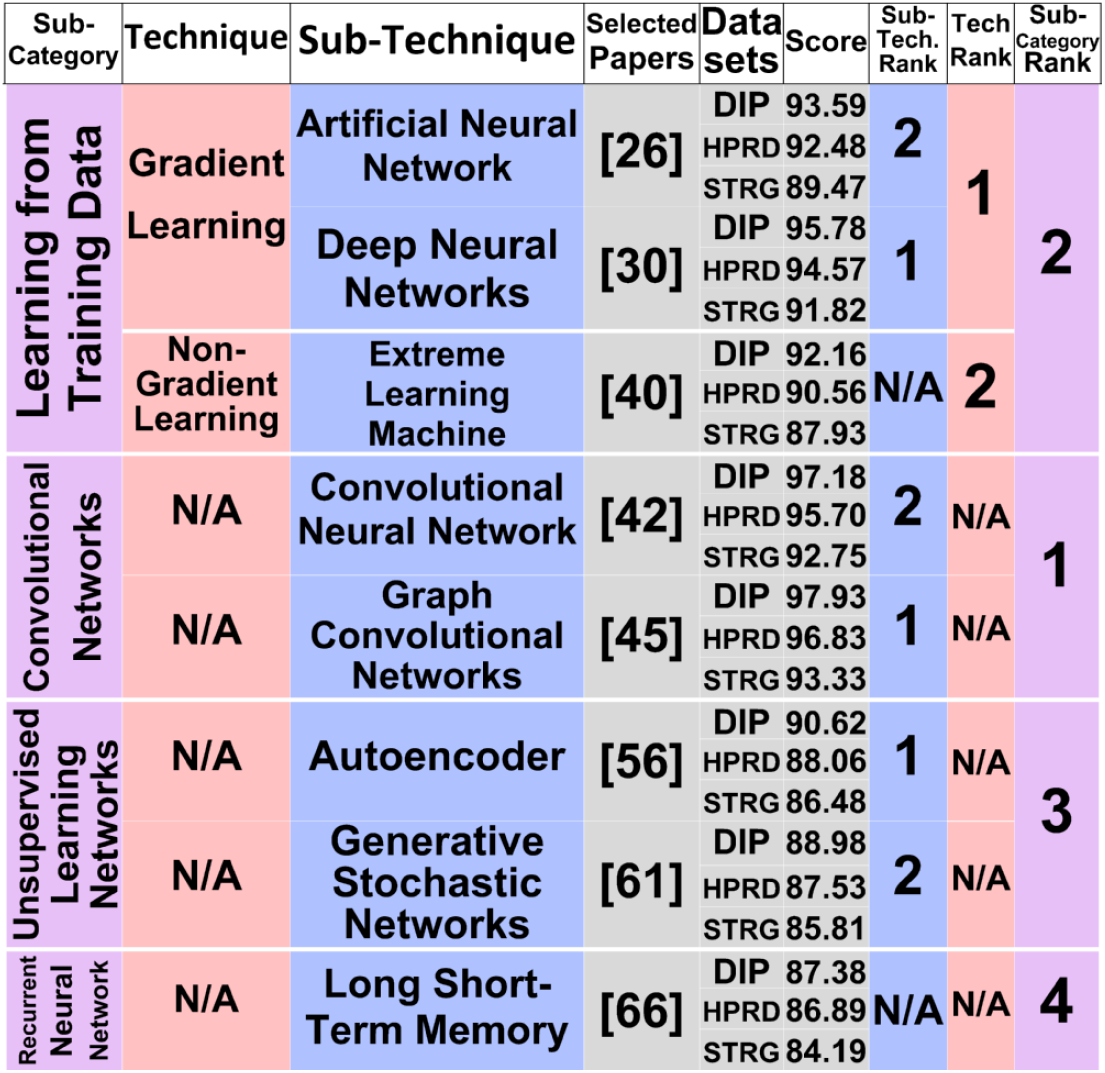
<u>Specificity</u> scores of the selected <u>Deep Learning algorithms</u>. The table also shows the following: the ranking of the sub-techniques that belong to the same overall technique, the ranking of the various techniques that belong to the same sub-category, and the ranking of the various sub-categories that belong to the same category

**Table 6.**
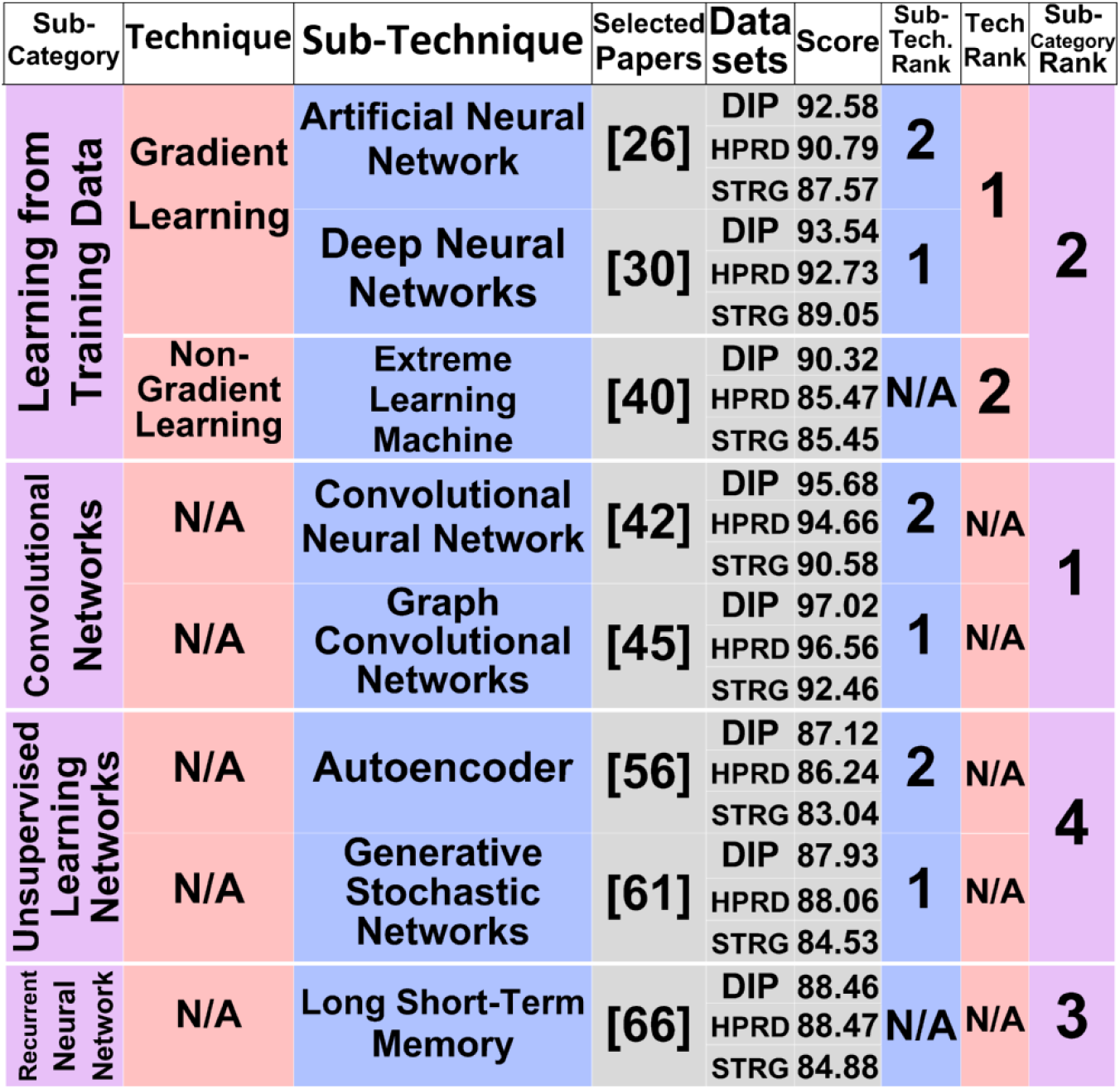
<u>Precision</u> scores of the selected <u>Deep Learning algorithms</u>. The table also shows the following: the ranking of the sub-techniques that belong to the same overall technique, the ranking of the various techniques that belong to the same sub-category, and the ranking of the various sub-categories that belong to the same category

**Table 7.**
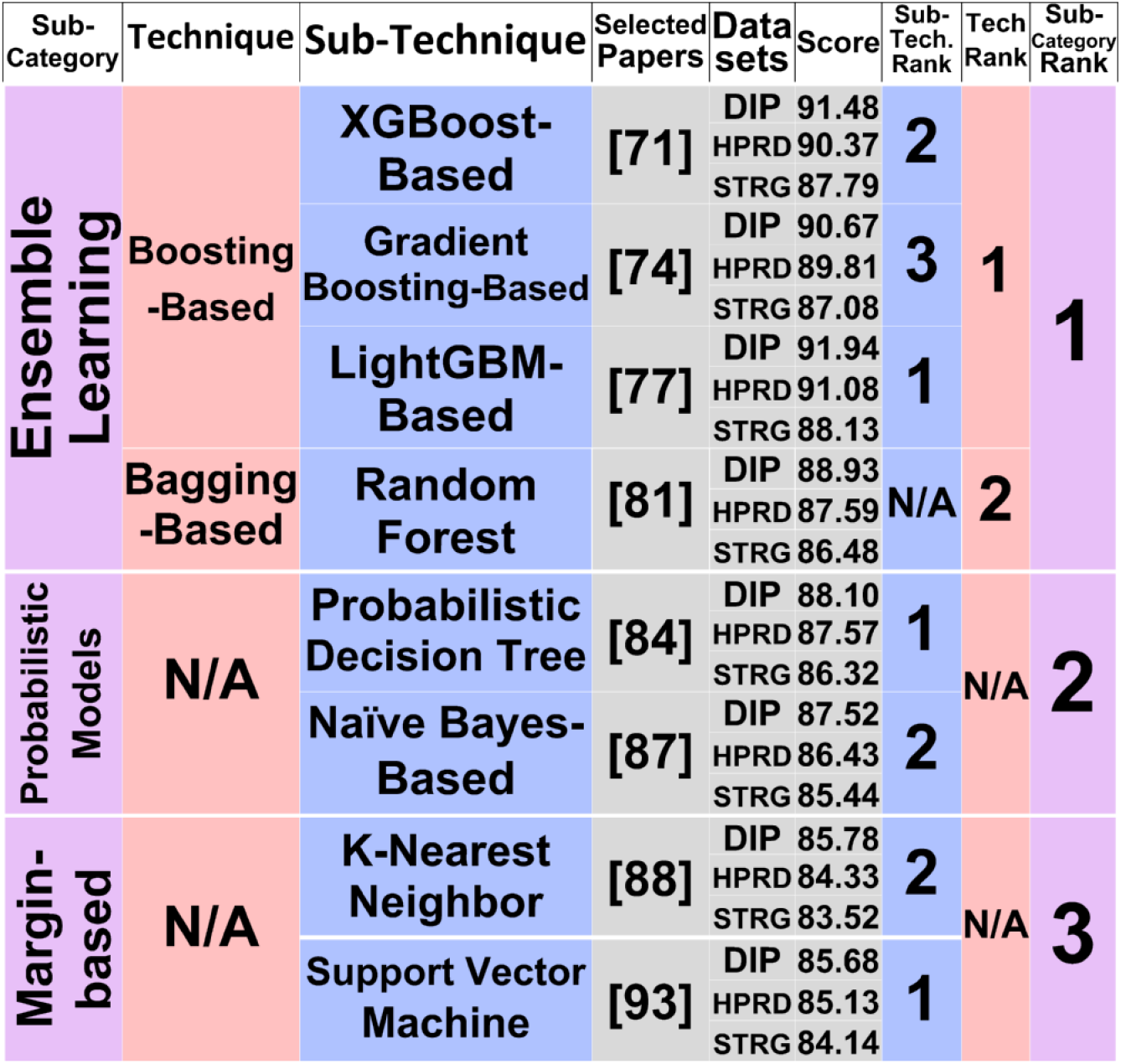
<u>Sensitivity</u> scores of the selected <u>Traditional Machine Learning algorithms</u>. The table also shows the following: the ranking of the sub-techniques that belong to the same overall technique, the ranking of the various techniques that belong to the same sub-category, and the ranking of the various sub-categories that belong to the same category

**Table 8.**
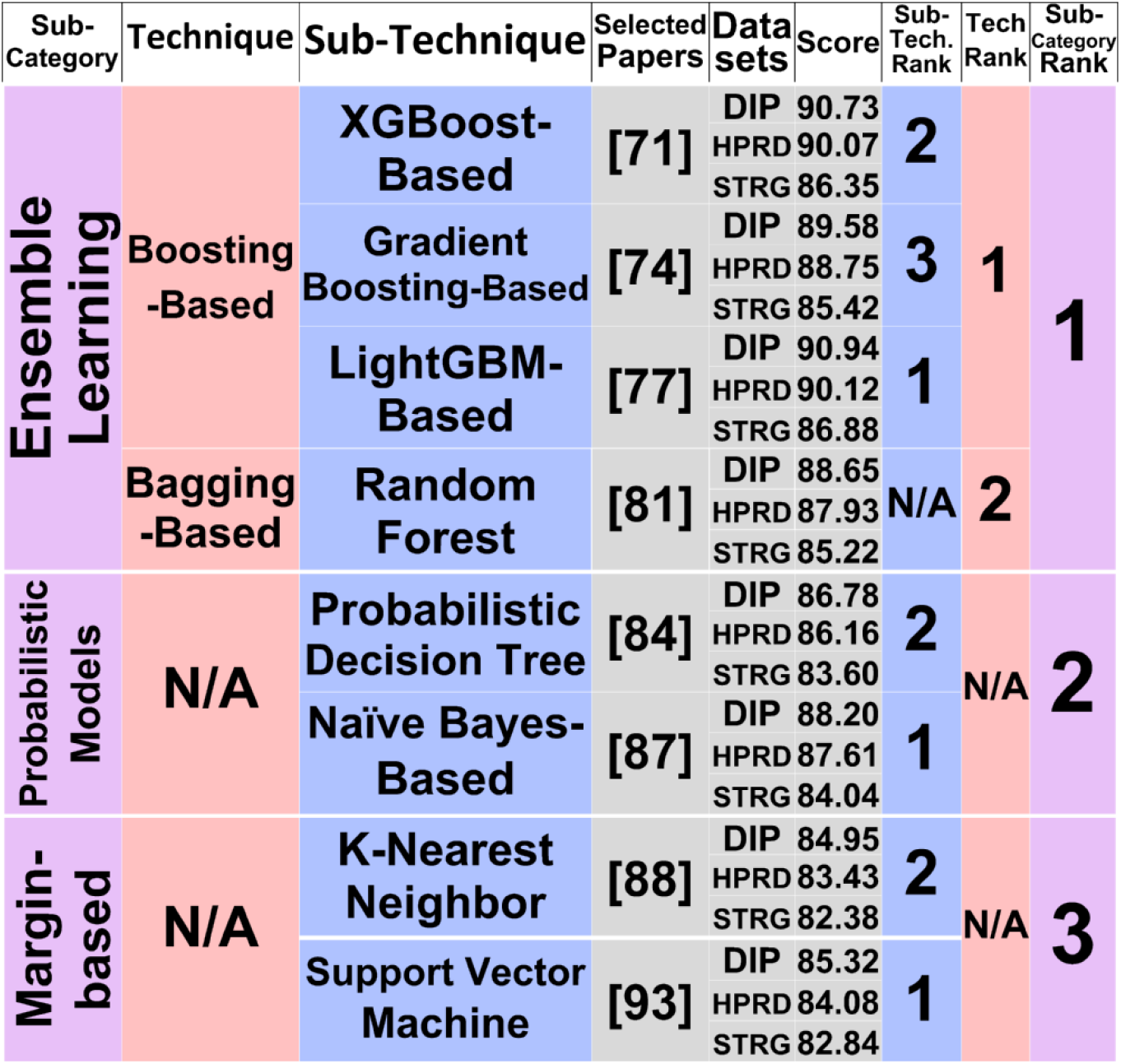
<u>Specificity</u> scores of the selected <u>Traditional Machine Learning algorithms</u>. The table also shows the following: the ranking of the sub-techniques that belong to the same overall technique, the ranking of the various techniques that belong to the same sub-category, and the ranking of the various sub-categories that belong to the same category

**Table 9.**
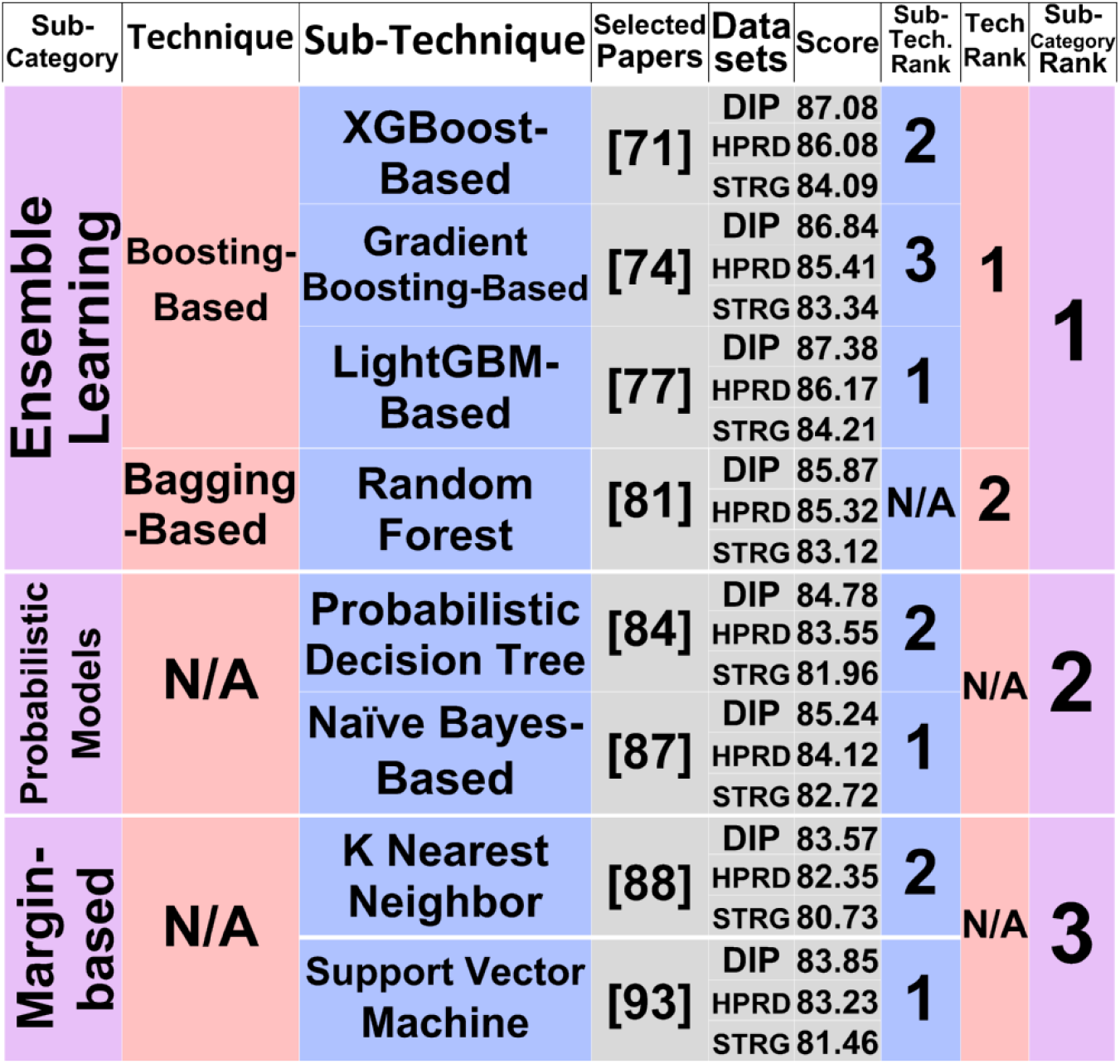
<u>Precision</u> scores of the selected <u>Traditional Machine Learning algorithms</u>. The table also shows the following: the ranking of the sub-techniques that belong to the same overall technique, the ranking of the various techniques that belong to the same sub-category, and the ranking of the various sub-categories that belong to the same category

We carried out an extensive search for codes that are publicly available and correspond to the algorithms we selected to illustrate their respective methods. Out of the algorithms chosen, we managed to obtain publicly accessible codes for seven of them. Below are the links to these codes, along with the papers where they have been reported on:

[30] https://github.com/xueleecs/SDNN-PPI

[45] https://github.com/lightaime/deep_gcns_torch and https://github.com/lightaime/deep_gcns

[66] http://kurata35.bio.kyutech.ac.jp/LSTM-PHV/

[71] https://github.com/SatyajitECE/DNN-XGB-for-PPI-Prediction

[77] https://github.com/QUST-AIBBDRC/LightGBM-PPI/

[88] https://sites.google.com/site/supplementaryfilestensymp19/downloaded-files

[93] http://www-tsujii.is.s.utokyo.ac.jp/satre/akane/

In relation to the remaining representative papers, we constructed our own versions using TensorFlow, following the procedures detailed by Sinaga and Yang [103]. These implementations were trained using the Adam optimizer, as recommended by Sinaga and Yang [103]. As noted in [104], TensorFlow’s APIs allow users to create their own algorithms. We conducted our development in Python 3.6 and used TensorFlow 2.10.0 as the underlying support for the models.

### 6.4 Ranking the Various Techniques, Sub-Techniques, Sub-Categories, and Categories

Tables 4-9 present the results of the experiments, as follows:

- *The scores of the deep learning algorithms:* Tables, 4, 5, and 6 show the scores of the sensitivity, Specificity, and precision scores, respectively. Each of these tables also shows the following: the ranking of the sub-techniques that belong to the same overall technique, the ranking of the various techniques that belong to the same sub-category, the ranking of the various sub-categories that belong to the same category.
- *The scores of the traditional machine learning algorithms:* Tables, 7, 8, and 9 show the scores of the sensitivity, Specificity, and precision scores, respectively. Each of these tables also shows the following: the ranking of the sub-techniques that belong to the same overall technique, the ranking of the various techniques that belong to the same sub-category, the ranking of the various sub-categories that belong to the same category.

### 6.5 Discussion of the Experimental Results

#### 6.5.1 Deep Learning-Based Techniques

##### 6.5.1.1 Artificial Neural Network (ANN)

The ANN model designed for PPI detection exhibited commendable predictive accuracy in the experimental findings. By utilizing convolutional layers to analyze sequence data, the model succeeded in recognizing spatial connections that might be overlooked by other techniques. In the context of PPI detection, this means the layers could capture local patterns and intricate structures within the sequence data, identifying relationships that are often beyond the reach of traditional analysis methods. Unlike simpler approaches that might rely on manual feature extraction, convolutional layers in the ANN model could recognize these spatial connections using filters and pooling layers, automatically learning the most relevant features.

Nevertheless, the approach was subject to certain constraints: (1) a tendency to overfit, though this was alleviated by dropout and regularization methods, (2) difficulties with less common proteins, a problem that was mitigated through data augmentation and the application of transfer learning from analogous tasks, (3) an imbalance of classes, which was addressed by oversampling the minority class and employing class weights during training to counteract misclassification of the less-represented class, (4) lack of interpretability, and (5) failure to accurately detect proteins with a limited presence in the dataset.

##### 6.5.1.2 Deep Neural Networks (DNN)

The experimental findings demonstrated that the method delivered remarkable performance, surpassing the majority of deep learning and conventional machine learning models. The convolutional layers appeared to adeptly identify spatial connections between amino acids. By integrating convolutional and recurrent layers, the model managed to discern both the spatial and sequential characteristics of proteins. The attention maps gave valuable insights into the principal residues participating in the interaction. These maps are essentially visualizations that demonstrate how the network is focusing its attention on different parts of the input, in this case, the amino acid residues involved in PPI. By highlighting the specific residues that play a crucial role in the interaction, the attention maps enabled a deeper understanding of the underlying biological mechanisms.

Nevertheless, the technique exhibited several constraints: (1) the model’s performance was subpar with certain protein classes, though this was partially alleviated by augmentation and adding features like secondary structure, (2) specific classes performed below expectations, and (3) overfitting occurred, which was mitigated through the use of dropout layers, meticulous cross-validation, and ensuring the robustness of results across various datasets.

##### 6.5.1.3 Extreme Learning Machine (ELM)

The outcomes from the ELM approach were notably encouraging. The model’s accuracy surpassed that of traditional machine learning techniques, and its training speed was a prominent aspect that differentiated it. Achieving results comparable to certain deep learning models, the ELM did so with only a fraction of the computational time required. The ELM model’s performance was consistent across various datasets, displaying not just high precision but also the capacity to generalize. Utilizing k-fold cross-validation and judiciously choosing the number of hidden neurons contributed to avoiding overfitting. The inherent simplicity of the ELM aided in optimization without compromising effectiveness.

##### 6.5.1.4 Convolutional Neural Network (CNN)

The CNN’s experimental outcomes ranked it second best among both traditional machine learning and deep learning models, emphasizing the power of deep learning in predicting intricate biological interactions. By utilizing a combination of oversampling and data augmentation methods, the model overcame the imbalance issue. The convolutional layers were fine-tuned to detect spatial features pertinent to PPI. Despite the model’s high computational demands, its interpretability was enhanced by visualizing the convolutional filters. The implementation of dropout layers along with regularization techniques played a key role in mitigating overfitting. Dropout is a regularization technique where during training, randomly selected neurons are ignored or “dropped out.” This means that they are effectively removed during a particular forward or backward pass. By doing so, the network becomes less reliant on specific neurons and is forced to learn more robust and generalized features.

##### 6.5.1.5 Graph Convolutional Networks (GCN)

The research findings indicated a substantial enhancement in the accuracy of PPIs using GCNs. In fact, this approach recorded the greatest accuracy, surpassing all other existing deep learning and machine learning techniques. By utilizing a graph-based representation, the complex spatial information within protein structures was effectively encapsulated, leading to a more insightful understanding of the relationships. Also, the consistent performance across various datasets attested to the method’s resilience and reliability. Features such as edge attributes and local receptive fields were instrumental in offering a transparent view of the interactions between proteins. However, there were certain sensitivities related to the selection of hyperparameters within the GCN, which required systematic tuning using methods like grid and random search to determine the optimal configuration. Though powerful, the method did present challenges in terms of computational demands, especially when dealing with expansive graphs.

##### 6.5.1.6 Generative Stochastic Networks (GSN)

Generally, the tests on the GSN for PPI detection demonstrated a moderate level of accuracy but encountered problems with the model’s convergence. The stochastic characteristics appeared to heighten the complexity. By experimenting with various hyperparameters and integrating regularization methods, the issue was alleviated to an extent. There was a fine equilibrium to maintain between grasping the intricacy of PPI and preserving the manageability of the model. The challenge is to find a fine equilibrium between these two competing needs. It involves carefully selectin features and representations that capture the essence of PPIs without overwhelming the model. Techniques like dimensionality reduction, regularization, feature engineering, and model selection come into play here. The model’s architecture must be designed in a way that it can learn from the available data efficiently, without becoming too cumbersome to train or interpret.

##### 6.5.1.7 Long Short-Term Memory (LSTM)

The experimental findings demonstrated that the LSTM method for PPI detection was reasonably accurate. Through identifying temporal dependencies in sequence data, the LSTM model yielded satisfactory outcomes, exhibiting potential for uncovering complex PPI patterns. To manage the imbalance in the dataset, a combined approach of oversampling the minority class and undersampling the majority class was used. The application of feature scaling to normalize the input sequences notably improved the model’s effectiveness. In certain cases of PPI detection, the precision of the LSTM model seemed to fall short, an issue that was linked to the LSTM’s hyperparameters. The efficiency of the LSTM technique in dealing with sequential data and its ability to grasp long-term dependencies rendered it apt for managing the intricacy and noise found in PPI data.

#### 6.5.2 Traditional Machine Learning Techniques

##### 6.5.2.1 eXtreme Gradient Boosting (XGBoost)

The experimental findings showed that using XGBoost for PPI detection resulted in remarkable outcomes, surpassing most conventional machine learning models. XGBoost’s gradient boosting framework was successful in significantly reducing classification errors, while the regularization term assisted in preventing overfitting. The combination of the gradient boosting technique and meticulous hyperparameter tuning contributed to exceptional accuracy. The ability to perform parallel computations was also a contributing factor. Its ability to scale also made it an appropriate choice for handling extensive datasets. When compared to deep learning models, XGBoost offered the benefits of quicker training and reduced computational complexity. XGBoost employs a gradient boosting framework that builds trees sequentially. Each tree corrects the errors made by the previous one, allowing for an efficient learning process. This can often lead to quicker convergence to a good solution compared to some deep learning models.

##### 6.5.2.2 Gradient Boosting (GBoosting)

The GBoosting model demonstrated outstanding efficacy in identifying PPI. Using grid search to pinpoint the best hyperparameters, the incorporation of L1 regularization markedly enhanced the model’s performance. In the protein sequences, GBoosting was able to discern the vital features that were most influential in predicting PPI. Notably, the composition of amino acids and the information regarding secondary structure proved to be highly consequential. The ability to interpret the GBoosting model was a beneficial feature. Nevertheless, the model did reveal some shortcomings when dealing with imbalanced data.

##### 6.5.2.3 Light Gradient Boosting Machine (LightGBM)

The experiment revealed that LightGBM excelled in predicting PPIs, achieving a performance that surpassed all other traditional machine learning models. Its efficiency and ability to scale were particularly prominent when handling large and skewed datasets. When compared to other machine learning methods, LightGBM demonstrated enhanced accuracy and decreased computational time, affirming its appropriateness for this task. Nonetheless, the model’s dependence on hyperparameter tuning rendered it susceptible to the selection of specific parameters.

##### 6.5.2.4 Random Forest

The Random Forest model displayed strong performance in the detection of PPI, reaching satisfactory accuracy levels. Using feature selection and hyperparameter optimization, it managed to identify complex interaction patterns. The assessment of feature importance revealed that both the amino acid composition and domain-domain interaction characteristics played vital roles in enhancing the predictive accuracy. This underscores the model’s potential as a valuable tool for predicting PPI. However, certain drawbacks were observed, including the challenge posed by the vast dimensionality of the feature space and occasional overfitting resulting from improper parameter configuration. These issues sometimes hindered the correct classification of specific interaction types, particularly when they were unevenly represented in the dataset. Also, the model’s training time was significantly more prolonged compared to other techniques.

##### 6.5.2.5 Probabilistic Decision Tree (PDT)

Utilizing PDT in PPI detection has yielded encouraging outcomes. The model managed to strike an acceptable equilibrium between sensitivity and specificity, and its probabilistic characteristics enabled it to encapsulate some of the innate complexity of PPIs. Despite these achievements, the issue of false positives was a significant concern. Also, PDTs demanded more computational resources than conventional decision trees, which could pose a constraint in the context of extensive PPI networks.

##### 6.5.2.6 K-Nearest Neighbor

The KNN model reached moderate success in identifying PPI, with its accuracy reflecting the intricate relationships in the data by weighing the significance of each neighbor. We observed a marked variation in performance when changing the number of neighbors, and using a weighted KNN further enhanced the results. The presence of imbalanced classes posed challenges for KNN, making it vital to either balance the dataset or employ metrics that are not as affected by imbalance.

##### 6.5.2.7 Support Vector Machine (SVM)

The SVM model displayed encouraging outcomes in identifying PPI, and the use of a Radial Basis Function (RBF) kernel appeared to be a practical option for the datasets. The model’s performance seemed to hinge on specific hyperparameters, such as the regularization parameter (C) and the kernel parameter (γ). The class imbalance problem was addressed using the Synthetic Minority Over-sampling Technique (SMOTE). By implementing cross-validation and an appropriate train-test division, the problem of overfitting was lessened.

## 7 Potential Future Perspectives on Machine Learning Techniques for Predicting PPI

### 7.1 Deep Learning-Based Techniques

#### 7.1.1 Artificial Neural Network (ANN)

Integrating diverse biological datasets and functional annotations can provide a comprehensive view of protein interactions, enriching the learning process. The development of new architectures and algorithms that specifically cater to the characteristics of PPI data could lead to more nuanced and insightful predictions. Combining ANNs with other machine learning methods in a hybrid approach may lead to more robust models that can capture the intricate patterns in PPI. Also, improving the interpretability of ANNs in the context of PPI will provide a better understanding of the underlying biological processes and assist in translating the findings into practical applications.

#### 7.1.2 Deep Neural Networks (DNN)

Improving data preprocessing techniques could lead to more accurate training data. This, coupled with innovative network architectures, might allow DNNs to better capture the underlying biology of PPIs. The integration of different data types, such as sequence information, structural information, and biological pathways, could provide a more comprehensive view. Leveraging domain knowledge and integrating it with model development might result in models that are more biologically relevant. In terms of efficiency, optimizing algorithms might lead to quicker training and inference times. This could make DNNs more accessible to a broader scientific community and enable real-time or near-real-time predictions.

#### 7.1.3 Extreme Learning Machine (ELM)

First, enhancing the initialization process of hidden layer weights can lead to more robust predictions. By finding optimal ways to initialize these weights, the convergence of the training process can be sped up, resulting in a more efficient model. Second, adopting advanced optimization techniques might enable better fine-tuning of the model’s parameters. By using techniques like genetic algorithms or simulated annealing, the ELM’s architecture could be tailored more precisely to the specific task of PPI prediction. Third, a hybrid approach that combines ELM with other machine learning techniques like deep learning could yield significant improvements. Such a hybrid model could draw from the strengths of each approach, achieving higher predictive accuracy and robustness. Finally, developing techniques for handling imbalanced data can improve the model’s performance. Since PPI data might be skewed towards more common interactions, adopting under-sampling or over-sampling strategies might provide a more balanced view of the interactions, leading to better predictions.

#### 7.1.4 Convolutional Neural Network (CNN)

Enhancing efficiency using transfer learning and multitask learning represents another frontier. By leveraging existing models or information from related domains, these approaches could significantly reduce the amount of labeled data needed, making the models more versatile and accessible. Collaboration and alignment with biological insights and domain expertise are vital to tailor CNNs to the specific and complex needs of PPI prediction. Bridging the gap between the fields of machine learning and biology can lead to more context-aware, specialized models that respond more effectively to the unique challenges of predicting PPI. Improved methods for addressing challenges like data imbalance and noise through advanced preprocessing, augmentation, or even synthetic data generation are also essential to achieving better training and validation.

#### 7.1.5 Graph Convolutional Networks (GCN)

First, scalability improvements, including computational efficiency, can enable the handling of larger and more complex biological graphs. Second, increased robustness and generalization through diverse data inclusion and techniques like transfer learning will make these models more adaptable across different biological scenarios. Third, the integration of multi-modal data types could offer a more comprehensive view of protein interactions, thereby enhancing predictive accuracy. Fourth, the development of methods to increase the interpretability of GCNs will foster greater understanding and trust in the predictions. Finally, enhanced representation learning that integrates various biological information can provide richer insights into protein interactions.

#### 7.1.6 Generative Stochastic Networks (GSN)

The potential future improvements include advancements in the architecture, training algorithms, or integration with other data sources to enhance the model’s accuracy and robustness. Leveraging more extensive and diverse datasets, incorporating domain knowledge, improving optimization techniques, and developing interpretable models are some areas where research could lead to substantial improvements. Integrating GSNs with other machine learning approaches and exploring novel techniques to handle the specific challenges of PPI prediction, such as dealing with imbalanced datasets and considering the spatial and temporal dynamics of protein interactions, could also contribute to future advancements in this field.

#### 7.1.7 Long Short-Term Memory (LSTM)

Enhancements may include developing specialized LSTM architectures tailored to the unique properties of protein data, which might include integrating LSTM with other deep learning models like convolutional neural networks. The application of advanced optimization methods, regularization techniques, and efficient hardware might significantly reduce training time and enhance performance. Integration of various types of biological data, such as gene expression and sequence information, could lead to more accurate predictions. Enhancing interpretability and visualization of the LSTM models can make them more user-friendly and scientifically robust. Ensuring that the models are robust to different data distributions and can generalize well to unseen data is crucial for their wider applicability. There may also be a focus on improving data representation and preprocessing to create more informative input data, possibly through advanced feature engineering that incorporates biological knowledge.

### 7.2 Traditional Machine Learning Techniques

#### 7.2.1 eXtreme Gradient Boosting (XGBoost)

Efficiency can be improved by optimizing computational processes, such as parallel processing and GPU acceleration, reducing both training and prediction times. Scalability improvements are needed to handle larger datasets and enable real-time predictions, possibly through distributed computing and cloud integration. Feature selection and engineering can also be refined to increase prediction accuracy. Improvements in robustness include better handling of imbalanced datasets, which are common in PPI predictions. This could involve developing novel sampling techniques or customized loss functions that focus on minority classes. Also, the integration of domain-specific knowledge, such as biological pathways, could enhance the model’s understanding of complex interactions, leading to more accurate predictions.

#### 7.2.2 Gradient Boosting (GBoosting)

One of the main areas of focus is enhancing the model’s efficiency and scalability, allowing it to handle larger datasets with higher dimensions. This might involve leveraging parallel processing or improving the algorithm itself. Another crucial aspect is to make the model more interpretable by developing visualization tools and techniques that can explain the model’s predictions more transparently. There is also room to optimize hyperparameters automatically, using methods like Grid Search or Random Search, which can lead to better prediction performance. Utilizing advanced feature engineering or automated feature selection may also lead to more robust models. Combining GBoosting with other models, such as deep learning, could create more powerful hybrid models capable of capturing complex patterns.

#### 7.2.3 Light Gradient Boosting Machine (LightGBM)

First, there is an opportunity to enhance the model’s accuracy by fine-tuning hyperparameters and incorporating advanced optimization techniques. The integration of new feature selection mechanisms can also aid in removing redundant and irrelevant features, which would lead to better prediction results. One promising area of research is the fusion of domain-specific knowledge into the model, which could be achieved through expert-informed features or a hybrid modeling approach that combines LightGBM with other machine learning algorithms tailored to PPI prediction. This would cater the model more specifically to the unique characteristics of PPI. Further, improvements in the computational efficiency of LightGBM can make it more applicable to large-scale PPI datasets, unlocking new avenues for research. Finally, an exploration of interpretability and the provision of insights into how the model is making predictions could increase trust and allow for more nuanced adjustments to the model’s behavior. This may include investigating how different features and parameters contribute to predictions.

#### 7.2.4 Random Forest

Improving the algorithm’s hyperparameter tuning is an area for enhancement. Automated and more sophisticated hyperparameter optimization techniques, like Bayesian Optimization, can lead to models that are better suited to the unique characteristics of PPI data. Model interpretability can also be addressed, as Random Forest models can be seen as “black boxes.” Introducing techniques for model explanation and interpretation could make the model’s predictions more understandable and trustworthy to domain experts. Also, the efficiency and scalability of the Random Forest algorithm could be improved for handling the large and high-dimensional data that are often associated with PPI prediction. Lastly, integrating Random Forest with other machine learning techniques, like deep learning, could lead to hybrid models that capitalize on the strengths of different methods.

#### 7.2.5 Probabilistic Decision Tree (PDT)

This involves enhancements in the areas of accuracy, efficiency, and interpretability. Techniques like incorporating additional biological knowledge, improving feature selection, utilizing more advanced probabilistic models, and leveraging machine learning advancements may contribute to the enhancement of PDTs. Also, integrating with other predictive methods and using more extensive and diverse datasets may lead to more reliable predictions.

#### 7.2.6 K-Nearest Neighbor

Improving feature selection techniques can make the algorithm more effective by reducing the dimensionality of the data and focusing on the most informative attributes. Optimizing distance metrics that quantify the similarity between proteins can help the algorithm make more accurate predictions. Integrating domain-specific knowledge, such as biological pathways or molecular structures, can make the model more robust and meaningful in the context of PPI. Finally, ensemble methods, where multiple KNN models are combined, can improve the overall performance by reducing the chance of overfitting and leveraging different aspects of the data. Combining these strategies could lead to a more efficient and accurate KNN model for predicting PPI in future research. However, the specific details.

#### 7.2.7 Support Vector Machine (SVM)

For model optimization, methods like hyperparameter tuning and kernel selection can enhance the predictive accuracy. There is also an ongoing effort to develop specialized algorithms that can efficiently handle the unique characteristics of PPI data, such as imbalanced datasets. Feature selection represents another critical area for improvement, where novel algorithms could be devised to identify and incorporate only the most relevant features into the model, thus improving the efficiency and performance. Scalability is yet another concern. As biological data grows and complexity, future implementations of SVM for PPI prediction must be able to manage large-scale datasets without a significant loss in performance. Parallel computing and hardware acceleration might be key to this enhancement. Furthermore, the integration of SVM with other machine learning and statistical techniques could potentially lead to hybrid models that capitalize on the strengths of various approaches. This may open new avenues for predicting PPI with even higher accuracy and robustness. Finally, the application of deep learning techniques in conjunction with SVM may further improve PPI predictions, allowing the model to learn more complex patterns and representations from the data.

## 8 Conclusions

This survey paper offers an all-encompassing analysis of machine learning methods applied to PPI detection. We explore the pros and cons of PPI, aiming to furnish a foundation for upcoming studies in this field. Through both empirical and experimental investigations, we present a detailed examination of the latest algorithms. We also propose a new taxonomy, grounded in methodology, that ranks algorithms hierarchically, including comprehensive categories and particular methods. This classification system enables a thorough and orderly way to categorize algorithms, boosting researchers’ comprehension of the algorithms’ connections and methods. By leveraging this taxonomy, researchers can perform accurate assessments and contrasts of algorithms, obtaining a richer understanding of their attributes and deficiencies. The suggested taxonomy also acts as a roadmap for future investigations, steering the creation and appraisal of novel algorithms.

The survey doesn’t only furnish an in-depth classification scheme for machine learning PPI algorithms but also includes empirical and experimental assessments to gauge their efficacy. Our empirical investigation appraises machine learning techniques for identifying PPI according to four criteria. Through experimental scrutiny, we juxtapose and rank diverse algorithmic classes and methods, comprising those that use identical sub-techniques, varying sub-techniques within the same technique, diverse techniques within the same sub-category, and various sub-categories within a single category. The methodological taxonomy, combined with empirical evaluations and experimental contrasts, cumulatively augments researchers’ insight into existing machine learning algorithms for PPI identification. This enables them to make educated choices when tackling their research inquiries or challenges. In the following section, we highlight the primary findings from our experimental results. Below, we present the main discoveries from our experimental outcomes:

- The deep learning techniques yielded the best results:

- *Convolutional Neural Network (CNN):*The imbalance issue in the model was tackled by using a mix of oversampling and data augmentation strategies. The convolutional layers were carefully adjusted to recognize spatial features related to PPI. While the model required significant computational resources, its interpretability was improved by visualizing the convolutional filters. The inclusion of dropout layers and regularization methods was vital in preventing overfitting.
- *Graph Convolutional Networks (GCN):* By employing graph-based depiction, the intricate spatial information found in protein structures was adeptly captured, resulting in a more profound comprehension of the connections. The method’s consistent success across different datasets demonstrated its robustness and dependability. The utilization of features like edge attributes and local receptive fields provided a clear perspective on the interactions among proteins.
- The traditional machine learning techniques yielded the best results:

- *Light Gradient Boosting Machine (LightGBM)*: This method’s efficiency in scaling and its pronounced performance in dealing with large and skewed datasets set it apart. LightGBM outperformed other machine learning techniques by delivering improved accuracy and reduced processing time, thereby validating its suitability for the task.
- *eXtreme Gradient Boosting (XGBoost):* Utilizing XGBoost’s gradient boosting framework led to a substantial reduction in classification errors, and the regularization term helped in averting overfitting. The integration of gradient boosting and careful hyperparameter adjustment resulted in extraordinary accuracy. The capacity for parallel computations further added to its effectiveness. XGBoost’s scalability made it an apt selection for processing extensive datasets. In comparison to deep learning models, XGBoost provided the advantages of faster training and decreased computational intricacy.

